# Chemical targeting of the ATXN1 aa99-163 interaction site suppresses polyQ-expanded protein dimerization

**DOI:** 10.1101/2025.03.17.643445

**Authors:** Ioannis Gkekas, Katerina Pliatsika, Stelios Mylonas, Sotiris Katsamakas, Apostolos Axenopoulos, Simona Kostova, Erich E. Wanker, Dimitra Hadjipavlou-Litina, Konstantinos Xanthopoulos, Petros Daras, Spyros Petrakis

**Author notes:** These authors contributed equally (Ioannis Gkekas, Katerina Pliatsika). Corresponding Author Spyros Petrakis.

## Abstract

Spinocerebellar ataxia type 1 (SCA1) is a neurodegenerative disease caused by the expansion of a polyglutamine (polyQ) tract in the ATXN1 protein. This expansion is thought to be responsible for the gradual aggregation of the mutant protein, which is associated with increased cytotoxicity and neuronal cell death. Apart from the polyQ tract, other domains in ATXN1 are also involved in the initial events of protein aggregation such as a dimerization domain that promotes protein oligomerization. ATXN1 interacts with various proteins; among them, MED15 that significantly enhances the aggregation of the polyQ-expanded protein. Therefore, we set to identify the interaction site between ATXN1 and MED15 and assess whether its chemical targeting would affect polyQ protein aggregation. First, we predicted the structure of ATXN1 and MED15 and simulated their interaction. We experimentally validated that amino acids (aa) 99-163 of ATXN1 and aa548-665 of MED15 are critical for this protein-protein interaction (PPI). We also show that the aa99-163 domain in ATXN1 is involved in the dimerization of the mutant isoform. Targeting this domain with a chemical compound identified through virtual screening (Chembridge ID: 5755483) inhibited both the interaction of ATXN1 with MED15 and the dimerization of polyQ-expanded ATXN1. These results strengthen our assumption that the aa99-163 domain of ATXN1 may be involved in polyQ protein aggregation and highlight compound 5755483 as a potent first-in-class therapeutic agent for SCA1.

## Introduction

Spinocerebellar ataxia type 1 (SCA1) is an autosomal dominant, progressive, and fatal neurodegenerative disease. It is caused by the expansion of CAG trinucleotide repeats in the *ATXN1* gene encoding for polyglutamine (polyQ) residues in the relevant ataxin-1 (ATXN1) protein. Mutant ATXN1 exhibits a characteristic propensity for the formation of intranuclear inclusion bodies (IIBs) within affected neurons of SCA1 patients ^1,2^. Although *ATXN1* is broadly expressed in the brain, cerebellar Purkinje cells display selective neurodegeneration, which is associated with the ataxic phenotype in SCA1 ^3^. However, in recent years, it has become evident that other brain regions such as the brainstem, the cerebral cortex and the striatum are also affected in SCA1 ^4,5^.

ATXN1 contains multiple domains, each one playing a crucial role in diverse subcellular functions. contains multiple domains, each one playing a crucial role in diverse subcellular functions. Located at the N-terminal region, the polyQ tract (aa197-225) is necessary for disease development ^6,7^. Beyond the polyQ tract, other regions significantly contribute to the aggregation process. For example, the AXH domain promotes oligomerization, contributing to the aggregation of mutant ATXN1. Computational analysis indicates that deletion or replacement of the AXH domain reduces the aggregation propensity of ATXN1 ^8,9^. Additionally, both the self-association region (SAR, residues 494-604) and the C-terminal region (residues 690-816) are implicated in the self-interaction and aggregation of ATXN1. Moreover, the nuclear localization signal (NLS, residues 794-797) is a major determinant of nucleocytoplasmic shuttling of ATXN1 and mutations in NLS abolish toxicity in a transgenic model of SCA1 ^10^. These findings indicate that various domains within ATXN1 may affect its aggregation and potentially, disease progression.

ATXN1 is involved in numerous protein-protein interactions (PPIs). Partners include the transcription factors Senseless/Gfi-1 and Sp1, the mediator of retinoid and thyroid hormone receptors SMRT/SMRTER, the polyQ binding protein-1 (PQBP1), the RNA splicing factor U2AF65, the protein kinase A (PKA), and the transcriptional activator RAR-related orphan receptor alpha (RORα) ^11–15^. The interaction between ATXN1 and the transcriptional repressor Capicua (CIC) has been extensively studied ^16^. This interaction affects cerebellar Purkinje cell pathogenesis and mutations of key residues which suppress the interaction also reduce toxicity in a Purkinje cell-specific SCA1 mouse model ^17^. Furthermore, interactors may modulate the aggregation and proteotoxicity of the mutant ATXN1(Q82) protein. Notably, MED15, a subunit of the Mediator complex ^18^ was previously shown to enhance ATXN1 cytotoxicity and protein aggregation ^19^ potentially through coiled-coil (CC) interactions ^20^.

In this study, we aimed to identify the PPI site between ATXN1 and MED15 and investigate its involvement in the aggregation of polyQ-expanded ATXN1. Using computational and experimental methods, we show that ATXN1 interacts with MED15 through an N-terminal domain (aa99-163) located upstream of the polyQ region. This domain is also important for ATXN1 dimerization. Its targeting with a chemical compound predicted through virtual screening significantly suppresses the ATXN1 interaction with MED15 and the dimerization of the polyQ-expanded isoform.

## Results

### The aggregation of ATXN1(Q82) is affected by MED15

Experimental evidence indicates that various domains within polyQ-expanded ATXN1 may affect its aggregation. For example, the AXH domain (residues 562-693) is essential for its dimerization, contributing to SCA1 pathology ^8,9^. Therefore, we first assessed the effect of this domain in ATXN1 protein dimerization. A cDNA clone of ATXN1 lacking the AXH domain (ΔAXH) was generated and used for the construction of mCitrine-and NL-tagged LuTHy expression vectors ^21^. Compared to full-length (FL) ATXN1 which efficiently formed protein dimers, deletion of the AXH domain moderately suppressed protein dimerization of both the Q30 and Q82 isoforms (36% and 27%, respectively) in LuTHy assays (Supplementary Figure 1A-B). These results indicate that the AXH domain weakly affects ATXN1 dimerization, the initial step in the polyQ-mediated aggregation process in SCA1. They also suggest that other domains in ATXN1 are involved in its dimerization and aggregation.

We previously showed that the CC-rich protein MED15 interacts with the mutant isoform of ATXN1 and enhances its pathogenicity. This effect may be due to the expansion of CC regions adjacent to the N-terminus of the polyQ domain ^20^. MED15 induces ATXN1(Q82) aggregation and co-localizes with polyQ IIBs in neuroblastoma cells ^19^. Here, we studied the impact of MED15 in a primary cell model of ATXN1(Q82) protein aggregation with no measurable cytotoxicity ^22^. *mCherry-MED15* was stably overexpressed in MSCs inducibly co-expressing *YFP-ATXN1(Q82)* and production of recombinant proteins was validated by SDS-PAGE and immunoblotting. YFP-ATXN1(Q82) was detected at the expected molecular weight of 130 kDa, while the mCherry-MED15 protein was detectable at the appropriate molecular weight (115 kDa) only in Tet-On YFP-ATXN1(Q82) + mCherry-MED15 cells (Figure 1A). Tet-On YFP-ATXN1(Q82) + mCherry-MED15 cells also accumulated insoluble polyQ IIBs, as confirmed by a filter retardation assay (Figure 1B). Next, we investigated the morphology of YFP-ATXN1(Q82) IIBs in the presence or absence of mCherry-MED15. To this end, MSCs were induced to produce the polyQ-expanded ATXN1 and imaged with fluorescence microscopy 2, 5 and 10 days post-induction. Distinct morphological differences in ATXN1 IIBs were observed. At day 2, IIBs in cells co-expressing both transgenes were irregular in shape compared to the spherical IIBs in MSCs expressing only *YFP-ATXN1*(Q82). By day 10, IIBs seemed to fuse and grow in size (Figure 1C-D). Immunoblots of cell extracts at day 5 or day 10 indicated that the 130 kDa band corresponding to YFP-ATXN1(Q82) partially shifted towards a higher molecular weight band only in cells co-expressing *mCherry-MED15* (Figure 1E). These observations suggest that the aggregation of mutant ATXN1 is directly affected by its interacting partner MED15, as described previously ^19^.

**Figure 1.**
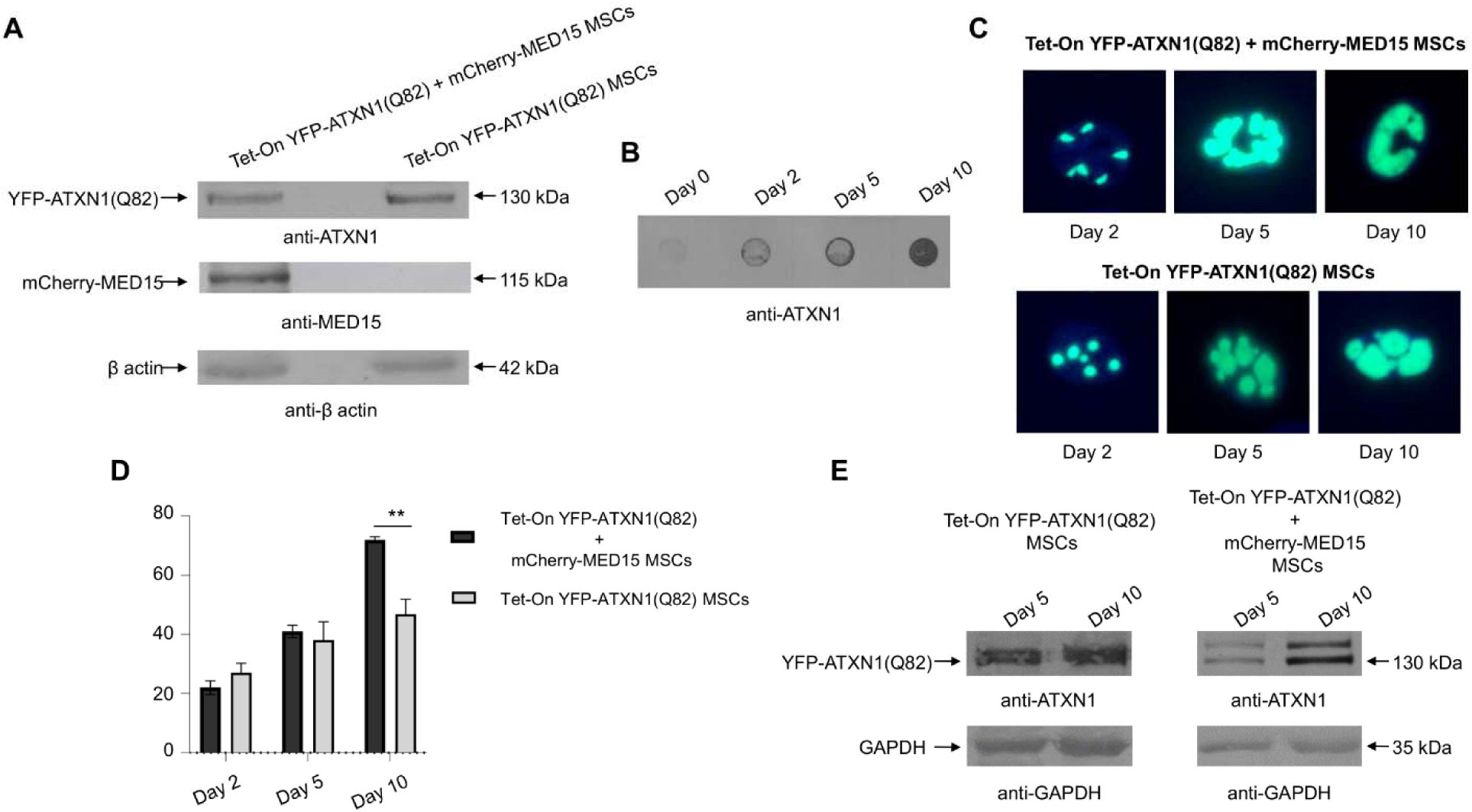
Effect of MED15 on the formation of ATXN1(Q82) IIBs. **(A)** Immunoblots for YFP-ATXN1(Q82) and mCherry-MED15 proteins in cell extracts of genetically-modified MSCs. β-actin was used as a loading control. **(B)** Filter retardation assay for the detection of insoluble YFP-ATXN1(Q82) IIBs in extracts from Tet-On YFP-ATXN1(Q82) + mCherry-MED15 MSCs. **(C)** Morphological analysis of YFP-ATXN1(Q82) IIBs at different time points in the presence (top row) or absence (bottom row) of mCherry-MED15 protein. Scale bar: 10 μM. **(D)** Bar graph showing the average size of YFP-ATXN1(Q82) inclusion bodies (IIBs) in MSCs, analyzed using the AggreCount software. The black bars represent cells producing only YFP-ATXN1(Q82), while the gray bars indicate cells co-producing mCherry-MED15. **(E)** Immunoblot for soluble YFP-ATXN1(Q82) protein at D5 and D10 post-induction. GAPDH was used as a loading control.

### ATXN1 interacts with MED15 through its aa99-163 domain

We hypothesized that MED15 exerts its effect on ATXN1(Q82) aggregation through a direct PPI. The interaction between ATXN1-MED15 has been previously validated along with the presence of MED15 in polyQ IIBs ^19^. To identify potential interaction sites, we simulated the ATXN1-MED15 PPI *in-silico*. In the absence of crystallographic data, the I-TASSER software suite ^23,24^ was utilized for a predictive modeling of the 3D structures of FL wild-type ATXN1 and MED15. Given the flexibility of the polyQ region in ATXN1, which complicates structural determination, we also predicted the structure ATXN1^NT^ and ATXN1^CT^, corresponding to protein fragments upstream and downstream of the polyQ tract, respectively (Supplementary Figure 2). Then, predicted structures of ATXN1 (ATXN1 FL, ATXN1^NT^ or ATXN1^CT^) were individually docked with two models of MED15 having the highest C-score. The most promising docking complexes were selected based on the score and reproducibility of docking solutions. Following docking simulations, a list of potential interaction sites between MED15 and ATXN1, ATXN1^NT^ or ATXN1^CT^ was generated. For example, the ATXN1 aa99-163 was identified as an interaction site between ATXN1^NT^ and MED15 FL (Supplementary Table 1).

The modeling capacity of protein prediction tools, including the recently developed AlphaFold, is strongly influenced by the presence of high-complexity domains in simulated proteins. This increases the possibility of low-confidence predictions ^25^. Thus, we sought to assess whether ATXN1 indeed interacts with MED15 through the predicted PPI sites. First, a LuTHy assay was established to quantify the already known ATXN1-MED15 PPI. Expression clones of mCitrine-ATXN1 and NL-MED15 were generated and used for the transfection of HEK293T cells. Control experiments were conducted using mCitrine-NL or combinations of mCitrine-ATXN1/NL and mCitrine/NL-MED15 plasmids. BRET measurements of control PA-NL were performed to correct for donor luminescence bleed-through. Cells producing mCitrine-ATXN1/NL-MED15 recombinant proteins exhibited a significantly higher BRET ratio compared to control interactions, providing a reproducible readout to quantify the ATXN1-MED15 PPI (Figure 2A and Supplementary Figure 3A-B).

**Figure 2.**
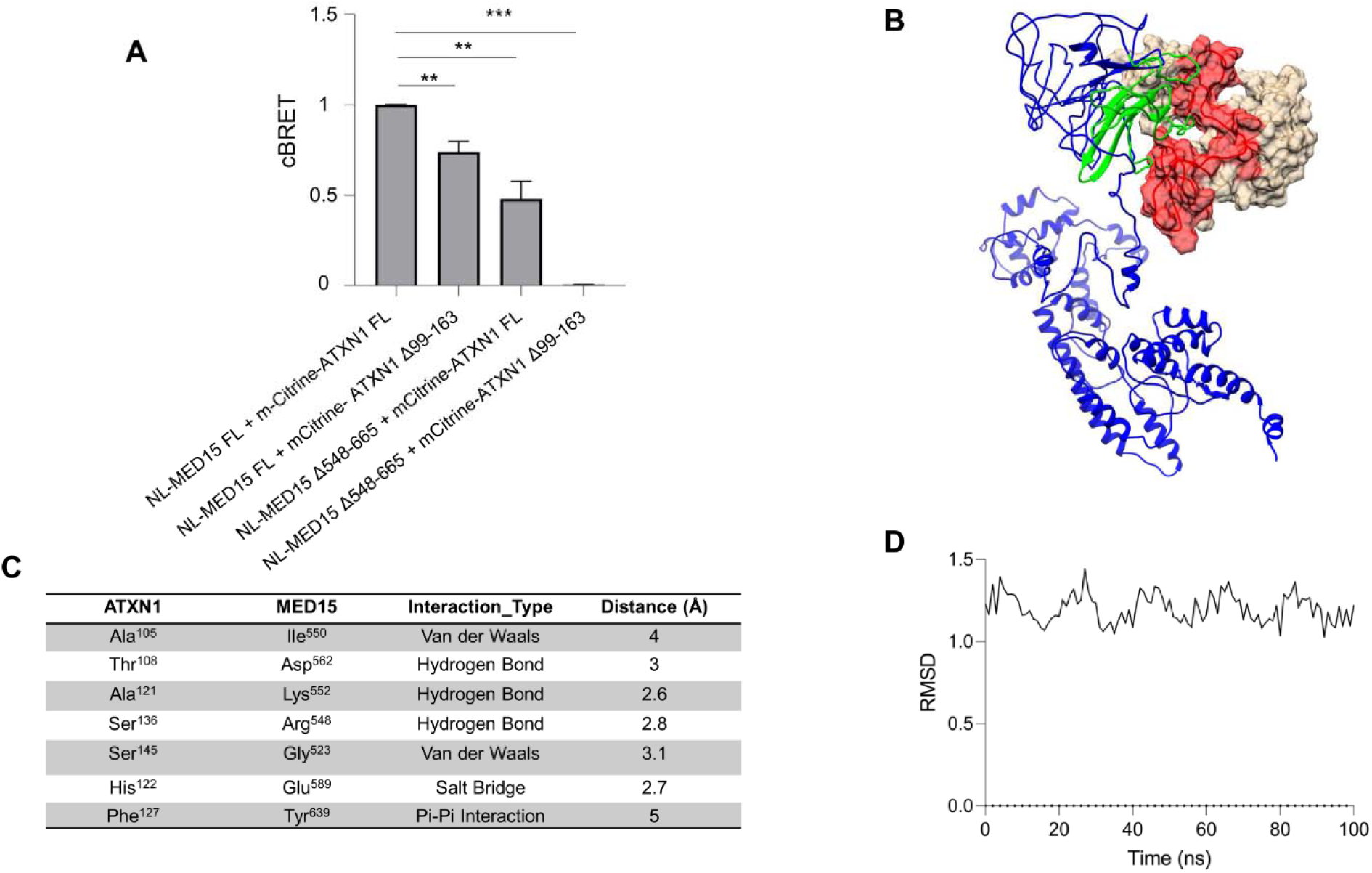
ATXN1 aa99-163 interacts with MED15 aa548-665. **(A)** LuTHy assay for the quantification of the interaction between full-length ATXN1 and MED15 or deletion-carrying proteins (ATXN1 Δ99-163 or MED15 Δ548-665). The simultaneous deletion of both domains from the interacting proteins abolishes ATXN1-MED15 PPI. Error bars denote mean ± SD (* p-value < 0.05, ** p < 0.01). **(B)** *In-silico* docking of ATXN1^NT^ and MED15 mediated through aa99-163 (red) in ATXN1 and aa548-665 in MED15 (green). The ATXN1^NT^ protein is shown in white while the MED15 protein is shown in blue, with the interaction domains highlighted. **(C)** Table summarizing the key residues mediating the ATXN1-MED15 PPI. It includes interacting residues in ATXN1 aa99-163 and MED15 aa548-665, along with the type of interaction and their respective distances (in Å). **(D)** RMSD of the ATXN1-MED15 complex over the simulation time.

Then, the predicted PPI sites were individually deleted from the full-length *ATXN1* and *MED15* genes. In total 5 deletion clones for *ATXN1* (Δ99-163, Δ198-248, Δ328-417, Δ547-691 and Δ493-540) and 5 deletion clones for *MED15* (Δ101-294, Δ195-294, Δ395-443, Δ548-665, Δ635-695) were generated and subsequently shuttled into LuTHy expression plasmids. The expression efficiency of these constructs was assessed by immunoblotting in extracts from transiently transfected HEK293T cells. Distinct protein bands corresponding to the respective ATXN1 and MED15 FL and deletion clones were detected at their expected molecular weight, confirming the successful production of deletion-carrying recombinant proteins (Supplementary Figure 3C-D). According to our hypothesis, deletion of the predicted PPI sites would result in a reduction or loss of the cBRET signal corresponding to the ATXN1-MED15 PPI. Therefore, PPIs between full-length ATXN1 and deletion clones of MED15 or deletion clones of ATXN1 and full-length MED15 were quantified using the LuTHy assay. Results indicated that deletion of aa99-163 or aa493-540 from ATXN1 resulted in a significant reduction of its PPI with MED15. Similarly, a reduction was observed when aa101-294, aa195-294 or aa548-665 region were deleted from MED15 (Supplementary Figure 4A). To exclude false positives, LuThy assays were repeated using combinations of hit ATXN1 and MED15 deletion clones. Notably, the simultaneous deletion of ATXN1 aa99-163 and MED15 aa548-665 resulted in a complete loss of the PPI (Figure 2A). In line with the experimental data, the computational prediction of the interaction between ATXN1^NT^ aa99-163 and MED15 aa548-665 is shown in Figure 2B. This data combined suggest that the interaction between ATXN1 and MED15 primarily occurs through this predicted PPI site.

To gain molecular-level information on the ATXN1-MED15 PPI, we performed a high-resolution docking analysis followed by MD simulations. These analyses highlighted key aa residues involved in the PPI, their interaction types and the interatomic distances stabilizing the protein complex. Molecular docking indicated the formation of three hydrogen bonds, one salt bridge, 2 Van Der Waals bonds and a Pi-Pi stacking interaction between key aa residues in ATXN1 and MED15 (Figure 2C). MD simulations also showed that these hydrogen bonds remained stable over time (Supplementary Figure 4B). Structural stability of the ATXN1-MED15 PPI was further analyzed through root mean square deviation (RMSD) and Radius of gyration (Rg) calculations, which indicate docking efficiency and distribution of atoms with respect to an axis of rotation, respectively. RMSD analysis confirmed a stable binding mode with a median RMSD of 0.12 nm and a minimal deviation over time (Figure 2D and Supplementary Table 2). Additionally, the Rg analysis suggested that the ATXN1-MED15 complex remained structurally compact with no significant expansion or collapse throughout the simulation. To further assess the stability and energetics of the ATXN1-MED15 PPI, we calculated the binding free energies over a 100-ns MS simulation using the Molecular Mechanics Poisson-Boltzmann Surface Area (MM-PBSA) approach. The computed binding free energy with a median of -10.4 kcal/mol, indicates a stable and energetically favorable interaction between ATXN1 and MED15 (Supplementary Table 2).

### The aa99-163 region is involved in ATXN1 dimerization

Sequences flanking the polyQ region are known to influence the dimerization and aggregation of various polyQ proteins ^26^. The aa99-163 domain of ATXN1 seems to be critical for its interaction with MED15. However, this 65-residue sequence, located upstream of the polyQ region may also mediate ATXN1 aggregation or other PPIs, including the self-interaction/homo-dimerization of polyQ-expanded ATXN1. To investigate this hypothesis, we deleted this domain from both wild-type ATXN1(Q30) and pathogenic ATXN1(Q82); then, we quantified the homo-dimerization of deletion-carrying ATXN1 proteins using the LuTHy assay. Analysis of the cBRET signal revealed that deletion of aa99-163 significantly suppressed the homo-dimerization of pathogenic ATXN1 compared to the full-length protein (Figure 3A). A similar suppression of homo-dimerization was also observed for ATXN1(Q30) (Supplementary Figure 5), which is known to self-associate and form soluble IIBs in overexpression models (Laidou et al., 2020). Donor saturation curves indicated significantly lower BRET ratios for Δ99-163 vs FL ATXN1(Q82) dimers suggesting a lower homo-dimerization affinity for the deletion-carrying mutant isoform (Figure 3B). These data suggest that the aa99-163 domain is not only critical for ATXN1-MED15 PPI but also important for ATXN1 self-interaction.

**Figure 3.**
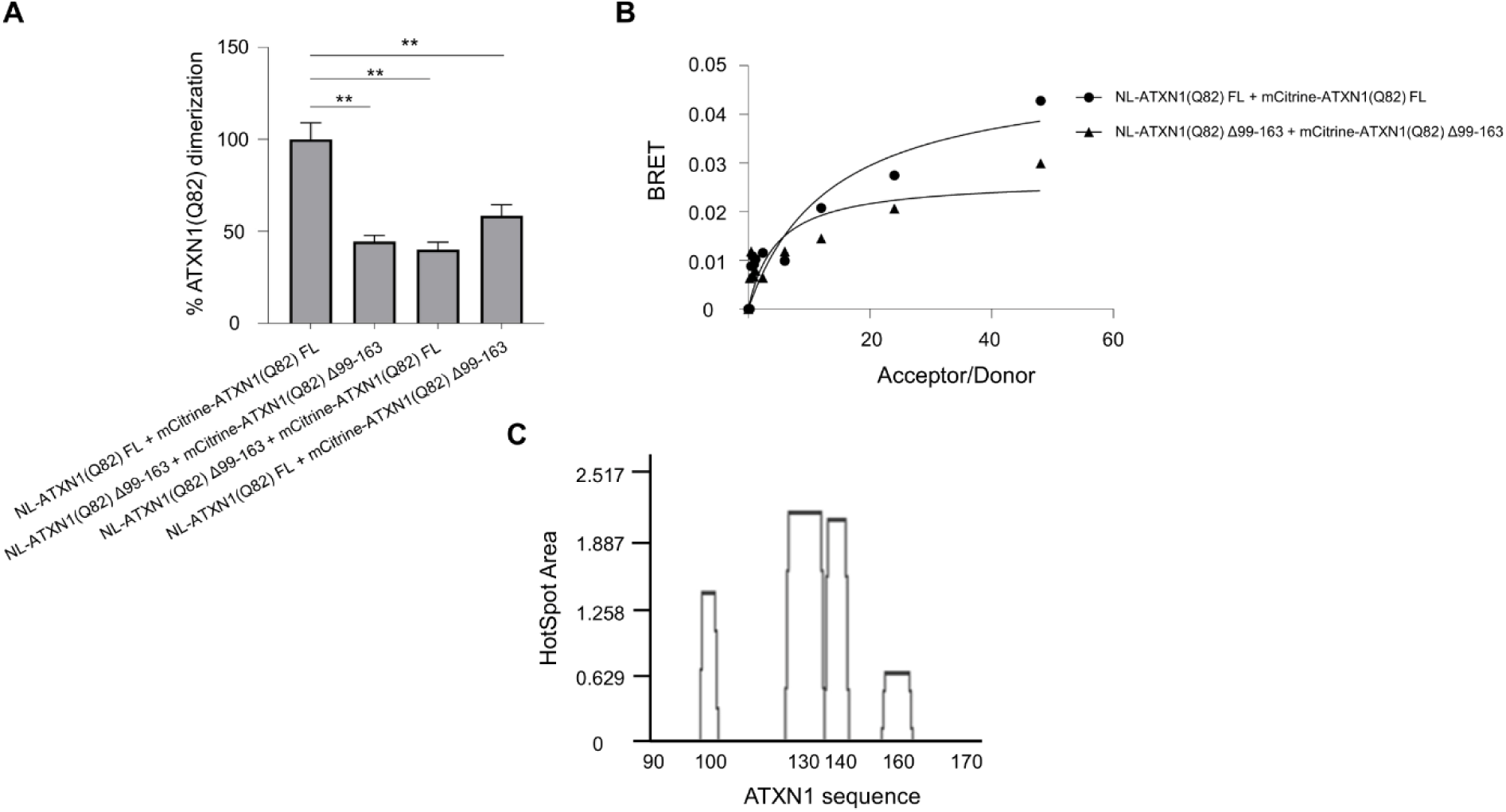
The aa99-163 domain is critical for the homo-dimerization of polyQ-expanded ATXN1. **(A)** Quantification of the homo-dimerization of full-length or Δ99-163 ATXN1(Q82) using the LuTHy assay. The bar graph shows various combinations of full-length or deletion-carrying ATXN1 proteins. Error bars denote mean ± SD (** p < 0.01). **(B)** Donor saturation assay indicating the effect of aa99-163 deletion in ATXN1(Q82) dimerization. **(C)** AGGRESCAN analysis indicating aggregation-prone regions at the N-terminal of ATXN1.

Given that dimerization is the initial step of protein aggregation, we further assessed the aggregation propensity of ATXN1 aa99-163. A sequence-based structural analysis wa performed using the AGGRESCAN algorithm. Key parameters associated with the aggregation profile (AP) of the polypeptide, including the average aggregation-propensity values per amino acid (a4v) and the HSA (Hot-Spot Area) for each aa residue were calculated. This computational analysis predicted an aggregation-prone aa stretch at residues 125-143 within ATXN1 aa99-163, suggesting that this domain has a high propensity for protein aggregation (Figure 3C and Supplementary Table 3).

### AI-based virtual screening for compounds binding to ATXN1 aa99-163

Since the aa99-163 domain is involved in ATXN1 interactions, we sought to identify chemical compounds that would bind to this domain and might be neuroprotective. To this end, we performed a virtual screening against this domain using the predicted ATXN1^NT^ structure which contains at least three compound binding pockets with high drugabbility probability (Supplementary Table 4). First, the commercially available Chembridge EXPRESS-Pick Stock library, comprising approximately 500,000 compounds was downloaded and pre-filtered according to a drug-like subset of pharmacokinetic (PK) properties to exclude non-favorable compounds from the process and minimize the load. Filtering included descriptors like Lipinski’s rule of 5. Furthermore, pan-assay interfering compounds (PAINS) and unwanted metabolites were removed and the Eli Lilly MedChem set of rules along with favorable PPI profile were applied. This pre-filtering step resulted in a reduced set of approximately 60,000 compounds for virtual screening. The pre-filtered compounds were progressed to virtual screening using an AI-assisted pipeline, which combined Smina docking results with a convolutional neural network (CNN)-based rescoring function. Our rescoring scheme assigned a low score to the majority of the screened compounds, while only a few of them were distinguished (Supplementary Figure 6A). We decided to qualify as potential hits the top-2% of the solutions, corresponding to 1,203 compounds with screening score higher than 0.901. These potential hits were further grouped into 214 clusters based on their structural similarity and only one compound per cluster was selected for the next phase. This final set of compounds was subjected to a secondary post-filtering step based on molecular modeling, properties and protein-ligand interactions. This rigorous selection process finally resulted in the identification of 24 hit compounds which are quite diverse in structure (Supplementary Figure 6B) and exhibited high binding scores to aa99-163 of ATXN1 (Supplementary Figure 6B and Supplementary Table 5). A schematic overview of the complete virtual screening pipeline is shown in Figure 4A.

**Figure 4.**
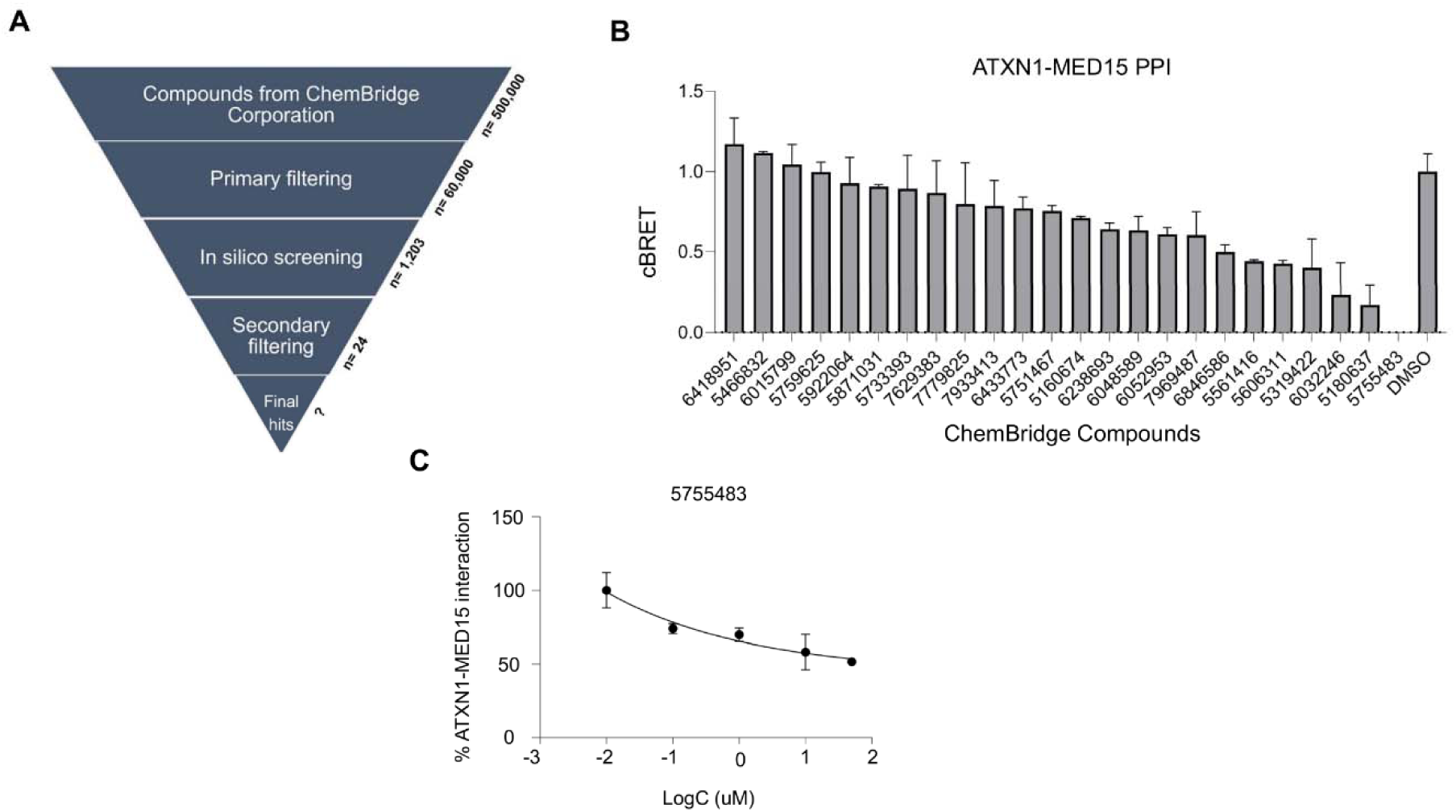
Screening for compounds suppressing ATXN1-MED15 PPI. **(A)** Workflow of the AI-based virtual screening. The screening process included compounds from ChemBridge Corporation, followed by primary filtering, computational docking to ATXN1 aa99-163 and secondary filtering for the selection of predicted hits. **(B)** Effect of computationally-predicted compounds on ATXN1-MED15 PPI using the LuTHy assay. Nine compounds significantly suppressed the interaction by at least 20% compared to the control DMSO-treatment, with compound 5755483 exhibiting the strongest inhibitory effect. **(C)** Concentration-response curve of compound 5755483 in the ATXN1-MED15 PPI. The x-axis represents the log concentration of the compound, while the y-axis shows the percentage of ATXN1-MED15 interaction.

### Compound 5755483 binds to ATXN1 aa99-163 and suppresses polyQ protein aggregation

The binding affinity of the 24 hit compounds for ATXN1 aa99-163 was evaluated in ligand-competition assays. First, we assessed their effect against ATXN1-MED15 PPI using the previously established LuTHy assay. Ιn principle, this interaction should be less strong compared to ATXN1(Q82) homo-dimerization which usually results in the formation of stable protein dimers and IIBs in cell-based overexpression assays ^19^. Transfected cells were treated for 48 hours with the highest non-cytotoxic concentration (Supplementary Figure 7) of each compound as indicated by the MTT assay, and those that reduced ATXN1-MED15 PPI by at least 20% were considered as positive hits. Based on this criterion, nine compounds (Chembridge ID: 5319422, 6433773, 5180637, 5606311, 5755483, 7969487, 6846586, 5751467, 6032246) significantly suppressed ATXN1-MED15 PPI compared to control samples treated with the solvent (DMSO). Notably, compound 5755483 (2’-({[4-(3-oxo-3-phenyl-1-propen-1-yl)phenyl]amino}carbonyl)-2-biphenylcarboxylic acid completely abolished the interaction (Figure 4B). Due to its structural characteristics the above compound might act as a Michael acceptor binding to several nucleophilic sites acting as PAINS. However pan-assay interfering compounds (PAINS) and unwanted metabolites were already removed and the Eli Lilly MedChem set of rules along with favorable PPI profile were applied. In order to remove any false positives, these nine compounds were further evaluated for their inhibitory effect in dose-response LuTHy assays. Indeed, six out of nine compounds tested, efficiently suppressed ATXN1-MED15 PPI in a dose-dependent manner (Supplementary Figures 8 and 9) with compound 5755483 demonstrating a potent effect (Figure 4C).

Finally, we investigated whether the binding of these compounds to ATXN1 aa99-163 also inhibit the dimerization of pathogenic ATXN1. Utilizing the previously established LuTHy assay for ATXN1(Q82) dimerization, we observed that only compound 5755483 (Figure 5A) suppressed mutant ATXN1 dimerization in a dose-dependent manner (Figure 5B). A similar effect was also observed in the dimerization of ATXN1(Q30) (Supplementary Figure 5B). To assess the stability of compound 5755483 binding on ATXN1 aa99-163, we performed a 100-ns MD simulation. This analysis indicated that the compound binds stably to ATXN1, with consistent interactions observed throughout the simulation. Below are given in details the amino acids implicated in the interactions and it is described the type of their observed interactions. More specifically, compound 5755483 forms a hydrogen bond with Ser^136^ with interaction frequency of 85% and mean distance of 0.30 nm (Figure 5C and Supplementary Figure 10A). Additionally, Thr^113^ and Pro^114^ form hydrophobic interactions with the compound showing an interaction frequency of 65% with mean distances of 0.2 nm and 0.24 nm, and binding free energies of -2.5 kcal/mol and -2.2 kcal/mol, respectively. These strong hydrophobic interactions contribute to the stabilization of the compound. Moreover, Tyr^135^ participated in a Pi-Pi stacking interaction with the compound, showing an interaction frequency of 67%, a mean distance of 0.35 nm and a binding free energy of -4.1 kcal/mol. The Pi-Pi stacking interaction with Tyr^135^ potentially enhances compound stabilization (Figure 5C and Supplementary Table 6). RMSD showed that the protein-compound complex remained relatively stable, with a distance of 0.2–0.4 nm over the simulation period. Additionally, root mean square fluctuation (RMSF) used to measure the flexibility of amino acid residues in a protein and Rg analysis indicate that the aa99-163 domain was less flexible in the presence of the compound suggesting that compound binding contributes to the structural stabilization of ATXN1 protein (Supplementary Figure 10B-D). No role for a methionine or a cysteine amino acid (the sulfur containing acids in a protein chain) was noticed

**Figure 5.**
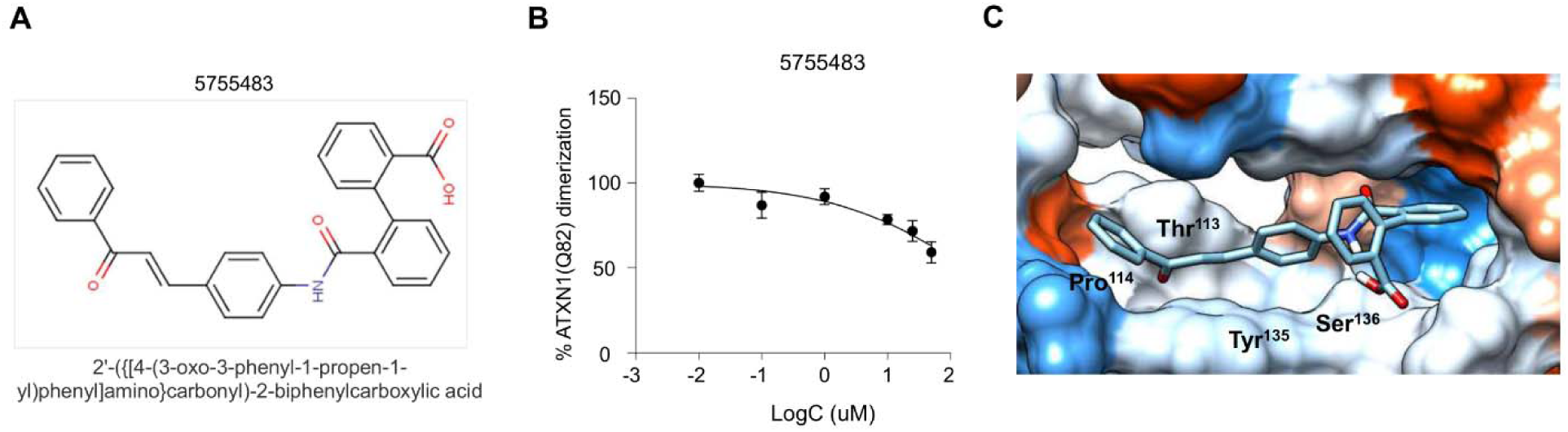
Compound 5755483 binds to ATXN1 and suppresses the homo-dimerization of pathogenic ATXN1. **(A)** Structure of the Chembridge compound 5755483. **(B)** Dose-dependent effect of compound 5755483 in the homo-dimerization of pathogenic ATXN1. **(C)** Docking simulation of compound 5755483 on the aa99-163 domain of ATXN1 identified the amino acids of ATXN1 that interact with the compound through hydrogen bonding, Pi-Pi stacking, or hydrophobic interactions.

## Discussion

### The role of PPIs in polyQ protein aggregation

PPIs involving polyQ proteins may play an important role in modulating their aggregation. Many of them may promote polyQ aggregation by facilitating the assembly of misfolded proteins into larger, more stable and insoluble structures. Conversely, other interactions may mitigate aggregation by enabling the clearance or degradation of these misfolded proteins. Understanding and characterizing these interactions is vital for identifying novel therapeutic targets in polyQ disorders. Compounds that would modulate these PPIs hold significant potential to reduce protein aggregation and alleviate the pathological effects associated with the presence of mutant polyQ proteins.

We have previously shown that MED15 interacts with ATXN1 and significantly promotes the aggregation propensity of its mutant isoform ^19^. This effect might be mediated by protein interactions promoting the inter-molecular assembly of polyQ-expanded ATXN1 into β sheet-rich fibrils ^20^. To investigate this hypothesis, we overexpressed MED15 in a previously established protein aggregation cell model of polyQ-expanded ATXN1. We observed that YFP-ATXN1(Q82) IIBs display a distinct morphology in the presence of MED15, suggesting that this protein may act as a linker between aggregation-prone molecules of mutant ATXN1. These results suggest that MED15 may directly facilitate polyQ aggregation.

We further focused on identifying a potential interaction site between the ATXN1-MED15 proteins. Using a computational approach, we found that this PPI may be facilitated by the ATXN1 aa99-163 domain, located at the N terminus of the protein. Computational approaches provide powerful predictions of protein structures and interactions but come with potential inaccuracies in modeling ^25^. We addressed this limitation by experimental validation. This dual approach enhances the reliability of our results and establishes a basis for the identification of critical interaction sites. While most known interactions of ATXN1 have been mapped onto the C terminus of the protein ^27–29^, our findings suggest an involvement of the N-terminal aa99–163 domain in mediating ATXN1 PPIs.

### The aa99-163 domain mediates ATXN1 homo-dimerization

Due to the absence of experimental data, the exact structure of ATXN1 remains elusive, hampering the design of efficient anti-aggregating approaches. Here, we show that deletion of the aa99-163 domain, upstream of the polyQ stretch, significantly suppressed the dimerization of pathogenic ATXN1, which may be considered as the first step in polyQ protein aggregation. Computational analysis using the AGGRESCAN software confirmed our hypothesis that this domain has a high aggregation propensity. Collectively, these findings suggest the diverse functionality of the ATXN1 aa99-163 domain, not only in mediating PPIs but also in ATXN1 homo-dimerization and potentially, aggregation.

Numerous studies indicate effects of protein domains other than the polyQ stretch in protein aggregation ^30–38^. These studies suggest that sequences flanking the polyQ tract significantly affect the aggregation process. For example, the addition of a 10-residue polyP sequence at the C-terminus of a polyQ peptide alters both its conformational properties and its aggregation kinetics. In Huntington’s disease (HD), the first seventeen amino acids at the N terminus (Nt17) of huntingtin influence its biochemical properties and the stability of polyQ aggregates, ultimately affecting disease progression. The initial phase of aggregation seems to start with the homo-dimerization of Nt17 and the formation of small, α-helix rich oligomers followed by the formation of fibrils also mediated by Nt17 ^39–46^. In SCA1, the AXH domain mediates ATXN1 interaction with the transcriptional repressor CIC. Disruption of the ATXN1-CIC complex ameliorates SCA1-like phenotypes in mouse models.

These examples highlight that chemical targeting of domains in polyQ proteins which mediate PPIs may represent a novel therapeutic strategy. Additionally, chemical compounds with high affinity for ATXN1 are largely missing. Using a combination of virtual screening and experimental validation we identified compound 5755483 (2’-({[4-(3-oxo-3-phenyl-1-propen-1-yl)phenyl]amino}carbonyl)-2-biphenylcarboxylic acid), which binds to ATXN1 aa99-163. Of particular significance is the binding of this compound to Ser^136^. Serine residues are known to participate in hydrogen bonding due to the presence of a hydroxyl group in their side chain, stabilizing protein-ligand interactions. Notably, compound 5755483 disrupted both ATXN1-MED15 PPI and the dimerization of pathogenic ATXN1. Its dual functionality highlights the therapeutic value of targeting specific interaction sites within ATXN1 to modulate polyQ protein aggregation.

## Conclusions

Together, our results indicate that the aa99-163 domain of ATXN1 is involved in a protein interaction with the aggregation-enhancer MED15 and the homo-dimerization of mutant ATXN1. Its chemical targeting with compound 5755483 suppresses polyQ-expanded ATXN1 dimerization. This compound may serve as a basis for chemical optimization and the development of derivatives with *in vivo* efficacy protecting from SCA1 neurodegeneration.

## Materials and Methods

### Generation of Tet-On YFP-ATXN1(Q82) + mCherry-MED15 MSCs

Mesenchymal stem cells (MSCs) were cultured in Dulbecco’s Modified Eagle Medium (DMEM, Biowest), supplemented with 10% fetal bovine serum (FBS, Biowest) and 1X penicillin-streptomycin (Biowest). Cells were maintained in a humidified incubator at 37°C with 5% v/v CO_2_. For the generation of Tet-On YFP-ATXN1(Q82) + mCherry-MED15 MSCs, naive MSCs were isolated and characterized as previously described ^47^. MSCs were seeded at a density of 2x10^5^ cells per well in a 6-well plate. Transfection was performed using Xfect reagent (Clontech) with a mixture of four plasmids: pT2-Tet/O2-YFP-ATXN1(Q82), pT2-mCherry-MED15, pT2-TetR-neoR, and pCMV(CAT)T7-SB100 transposon plasmids at a ratio of 4:3:2:1, respectively. Transfected cells were selected at day 7 post-transfection using 100 ug/mL G418 (InvivoGen). For induction of the *YFP-ATXN1(Q82)* transgene, doxycycline (Dox, 2 ug/mL, Sigma-Aldrich) was added to the culture medium. The *mCherry-MED15* transgene was constitutively expressed in genetically-modified MSCs.

### Fluorescence Microscopy

Cells were fixed with 4% v/v formaldehyde (Applichem) in PBS (Biowest) for 10 min and permeabilized for 10 min with 0.1% v/v Triton-X 100 (Sigma-Aldrich). For nuclei staining, cells were incubated with DAPI (Biotium) for 5 min at room temperature. Cells were observed in a ZOE Fluorescent cell imager equipped with three fluorescence channels and an integrated digital camera (Bio-Rad).

### Filter retardation assay

Extracts of MSCs treated in the presence or absence of Dox were mixed with an equal volume of a 4% w/v Sodium Dodecyl Sulfate (SDS) supplemented with 100 mM DTT and heated at 95°C for 10 min. Next, the samples were diluted with 100 uL 0.2% w/v SDS (Applichem) and filtered through a 0.2 μm cellulose acetate membrane (Whatman, Merck). SDS-resistant inclusions retained on the membrane were detected using the anti-ATXN1 SA4645 antibody (1:1,000 v/v) ^19^. The intensity of SDS-resistant inclusions was quantified using the ImageJ analysis software v1.54f. Statistical analysis was performed using the GraphPad Prism software v9 (San Diego, USA).

### Aggrecount

IIBs were quantified using AggreCount ^48^, an ImageJ macro developed for the unbiased analysis of protein aggregates. The macro utilizes a threshold-based segmentation approach to determine the optimal threshold for each image, ensuring accurate identification of IIBs. A minimum cut-off size of 5 µm² was applied to include only biologically relevant IIBs identified, counted, and measured for size. The analysis was conducted at single-cell resolution, with twenty cells per image analyzed across five images per experimental condition, resulting in a total of 100 cells per condition. Data compilation involved recording the number and size of IIBs in each cell, allowing for a comprehensive evaluation of their distribution and characteristics.

### Immunoblotting

Cells were lysed in RIPA buffer containing protease/phosphatase inhibitors (Thermo Fisher Scientific) and benzonase (Calbiochem-Novagen). Cell extracts were analyzed by SDS-PAGE electrophoresis followed by electrotransfer onto polyvinylidene difluoride (PVDF) membranes (Thermo Fisher Scientific). PVDF membranes were blocked with 5% w/v non-fat dry milk in PBST for 1 hour at RT and incubated with antibodies against ATXN1 (SA4645, 1:1,000 v/v dilution) ^19^, MED15 (H00051586-M02, Abnova, 1:1,000 v/v dilution), GAPDH (5174, Cell Signaling Technologies, 1:1,000 v/v) or β-actin (4970, Cell Signaling Technologies, 1:1,000 v/v dilution). After incubation with the appropriate secondary alkaline phosphatase-conjugated antibody, protein bands were visualized using NBT/BCIP (Applichem).

### Prediction of 3D protein structures

The 3D structures of full-length ATXN1 and MED15, as well as the N-terminal (ATXN1^NT^, aa 1-196) and C-terminal (ATXN1^CT^, aa 229-819) truncated forms of ATXN1, located upstream and downstream of the polyQ region, respectively, were predicted using the Iterative Threading ASSEmbly Refinement (I-TASSER) software ^23,24^. The resulting 3D models were evaluated based on their C-score, which represents the estimated quality of the model. The model of each ΑΤΧΝ1 protein (full-length ATXN1, N-terminal or C-terminal truncated fragments) with the highest C-score was selected. For MED15, two 3D models with a similar C-score were selected for docking experiments.

### Pocket druggability prediction

The presence and druggability of pockets was predicted in the PockDrug server using the fpocket algorithm. A ligand proximity threshold of 5.5 Å was applied to determine potential ligand-binding sites. Pockets were analyzed using a set of descriptors to evaluate their druggability potential, including volume hull which represents the total volume of the detected pocket, and hydrophobicity (Hydrophobic Kyte), measured using the Kyte-Doolittle scale to assess the hydrophobic character of the pocket. The proportion of polar and aromatic residues was also evaluated. The druggability of each pocket was assessed using the druggability probability score. A probability score greater than 0.5 indicates a druggable pocket ^49^.

### Protein docking simulation

For docking experiments, the Hex software was utilized ^50^. Hex is based on spherical polar Fourier (SPF) correlations to accelerate calculation of the interaction energy between local patches of the two proteins. This energy term comprises both surface shape and electrostatic charge, providing a comprehensive evaluation of potential binding interactions. Hex generates a number of solutions, representing docked positions of the two proteins, which are sorted based on their interaction scores. For each combination of ATXN1^NT^, ATXN1^CT^, and ATXN1 with the two MED15 models, approximately 20 different docking solutions were generated, sorted by their docking scores. The docking solutions comprised either specific amino acid residues or groups of residues from each protein that demonstrated potential binding interactions. To ensure robustness and specificity, single amino acids were excluded from the analysis, focusing on the interactions involving groups of amino acids. The most promising docking sites were selected based on their reproducibility across multiple docking solutions and their spatial localization within the ATXN1 and MED15 proteins. The reproducibility of docking sites was determined by assessing their frequency of occurrence in the generated docking solutions. Potential docking sites that were found in close spatial proximity were grouped into larger domains. Stretches of a minimum of 30 amino acids were arbitrarily considered as a protein domain.

### Generation of ATXN1 and MED15 entry clones lacking interaction sites

To generate ATXN1(Q30), ATXN1(Q82) and MED15 clones lacking interaction sites, a PCR-based approach was employed. Phosphorylated primers (Supplementary Table 7) were designed to hybridize immediately after the binding sites. PCR reactions (25 uL volume) contained 1 ng of template DNA, 0.3 mM of KAPA dNTP Mix, 0.3 µM final concentration of each phosphorylated primer and 0.5 U of KAPA HiFi DNA polymerase in 1X KAPA HiFi GC buffer (Roche). The PCR protocol consisted of 30 cycles, with the following profile: 20 s denaturation at 98°C, 15 s annealing at 60°C and 2.5 min extension at 72°C in an Eppendorf apparatus. PCR products were purified from agarose gels and re-circularized using T4 DNA ligase. Ligation reactions composed of 300 ng DNA fragment, 1 uL T4 DNA ligase (Thermo Fisher Scientific) and 1X T4 ligase buffer in a total volume of 20 uL. Following an overnight incubation at 16°C, each ligation reaction mixture was used for the transformation of Mach1 *E. coli* cells. Transformed cells were grown on agar plates supplemented with spectinomycin for MED15 deletion clones or kanamycin for ATXN1 deletion clones. Single colonies were selected and grown in LB medium supplemented with the respective antibiotic. Finally, plasmid DNA were purified from *E. coli* cells using the NucleoSpin Plasmid Kit, following the manufacturer’s instructions (Macherey-Nagel) and analyzed by BsrGI (NEB) restriction digestion. Deletion clones with the expected size were subjected to further verification of their identity via DNA sequencing.

### Generation of ATXN1 and MED15 expression clones

Full length ATXN1 and MED15 entry clones, as well as ATXN1 and MED15 entry clones with the desired deletions of binding sites were shuttled into pcDNA3.1-PA-mCitrine-GW or pcDNA3.1-myc-NL-GW Gateway destination vectors ^21^ Recombination was performed using LR clonase enzyme (Invitrogen). Reaction mixtures were incubated at 25°C and used for the transformation of Mach1 *E. coli* cells. Bacteria were grown on LB agar plates supplemented with 100 ug/mL Ampicillin. Plasmid extraction was carried out using the NucleoSpin Plasmid Kit. The identity of resulting plasmids was checked by BsrGI restriction digestion.

### Cytotoxicity assays

The cytotoxicity of the compounds was assessed using the MTT assay. HEK293T cells were cultured in 96-well plates at a seeding density of 4x10^4^ cells per well at 37°C in a 5% v/v CO_2_ atmosphere. The next day, cells were treated with various concentrations of the chemical compounds [dissolved and serially diluted in dimethyl sulfoxide (DMSO, Applichem) with concentrations ranging from 100 to 0.1 uM] or the solvent (DMSO) alone, serving as a control. The final concentration of DMSO in the culture medium was adjusted to 1% v/v. Following an appropriate incubation period of 48 hours, treated cells were exposed to 0.5 ug/mL of the MTT reagent (Applichem) and further incubated for 4 hours. Then, the MTT-containing medium was removed and the formazan crystals formed by viable cells were dissolved using DMSO. To quantify cell viability, the optical density (OD) of the formazan solution was measured at two wavelengths, 570 nm and 630 nm, using a SPARK plate reader (Tecan).

### LuTHy assay

PA-mCitrine-ATXN1/NL-MED15 were co-expressed in HEK293 cells (4x10^4^ cells per well). To assess the interaction between ATXN1 and MED15, BRET/cBRET signals were quantified 48 hours post-transfection. Luminescence emission was measured at short (370-480 nm) and long (520-570 nm) wavelengths after the addition of the appropriate substrate. For accurate analysis and correction of donor luminescence bleed-through, BRET measurements of the positive control PA-mCitrine-NL and the negative control PA-NL were performed. LuTHy assay^21^ was then performed for the quantification of PPIs between ATXN1 and deletion clones of MED15 or the opposite combination (full length MED15 and deletion clones of ATXN1). Luminescence measurements were used to quantify the interaction between full-length or deletion clones of ATXN1 and MED15. Protein production was estimated by total luminescence and fluorescence measurements. For donor saturation experiments, 0.5 ng of plasmid DNA encoding NL fusion proteins was co-transfected with increasing amounts (0.1–20 ng) of plasmid DNA encoding PA-mCit-tagged constructs. Measurements were performed 42 hours post-transfection and BRET ratios were determined to quantify interaction strength and efficiency. LuTHy assays were also performed in the presence of chemical compounds at various non-cytotoxic concentrations (8 – 48 hrs treatment). To ensure specificity and accuracy of the measurements, the non-specific/auto-fluorescence signal of each compound was calculated and subtracted from all relevant interactions. Then, the % inhibitory effect of the compounds was calculated by comparing the normalized cBRET signal of the treated cells to the solvent-treated control samples.

### Pre-filtering and profiling of compounds

The ChemBridge library, comprising 500,000 molecules in SDF formatted files, was downloaded from the Hit2Lead website (www.hit2lead.com). Prior to further analysis, this library was annotated to remove any present salts’ counter ions. This step was executed using OpenBabel v2.4.1 ^51^. Next, the library was submitted to the FAF-Drugs4 server ^52–56^ for stringent filtering based on drug-like properties, promiscuity, and PPIs considerations, employing a range of ADME-Tox descriptors such as XLOGP3, PKs, bioavailability, and adhering to Lilly Medchem relaxed rules ^57^. Compounds meeting all filtering criteria were compiled into SDF formatted files, forming the basis for subsequent virtual screening analyses.

### AI-based virtual screening

Following the pre-filtering stage, accepted molecules of SDF format were subjected to virtual screening. The virtual screening pipeline integrates a novel AI-based technique, where the docking outcome of Smina ^58^ is combined with an AI-driven rescoring function ^59^. This approach enhances the discernment of potential binding complexes with higher efficiency. Docking was performed on two different compound conformations resulting in fifty (50) docked poses per molecule conformation. The resulting protein-compound docked complexes were rescored using our custom AI-based rescoring function, a 3D neural network (3D-CNN) which has previously shown to outperform Smina in quantifying more effectively the protein-ligand binding mechanism ^59^. The top scoring molecules were further categorized into clusters based on their structural similarity, utilizing a 3D shape descriptor ^60^, and one molecule per cluster was finally qualified for further processing ^61^.

### Post-filtering of compounds

The remaining compounds were post-filtered based on three key criteria: molecular modeling, molecular properties, and the nature of protein-ligand interactions formed during docking simulations. This strategy ensured a rigorous evaluation process, encompassing both structural and functional aspects of the protein-ligand complexes.

### Molecular dynamics

Molecular dynamic (MD) simulations of protein-protein complexes and protein-inhibitor were performed using GROMACS 2024.4 ^62^. The CHARMM36 force field was used for all simulations. The system was resolved using a 1-nm dodecahedron water box, with chloride ions added to generate a charge-neutral system. The system was first minimized, followed by temperature and pressure equilibration under an NVT ensemble for 100 ps and then an NPT ensemble for an additional 100 ps. MD simulations were performed for 100 ns for each system at 300 K. Analysis of molecular dynamics was performed using GROMACS utilities and scripts in Python environment.

### Prediction of aggregation propensity

The aggregation propensity of the ATXN1(Q82) protein was analyzed using the AGGRESCAN algorithm ^63^. The peptide sequence of ATXN1(Q82) was submitted to the AGGRESCAN server in a FASTA format. Aggregation propensities of amino acids were calculated using the default settings. Key parameters such as the average aggregation-propensity values per amino acid (a4v) and the HSA (Hot-Spot Area) for each aa residue were calculated.

### Statistics and data fitting

All experimental assays were conducted in technical triplicates to ensure robustness and reproducibility of the results. The data are expressed as the mean ± standard deviation. To assess the potency of inhibitory compounds, the half-maximal inhibitory concentration (IC_50_) values were determined. Inhibition data were converted to % inhibition and fitted using a standard log inhibitor vs. normalized response model through nonlinear regression analysis in GraphPad Prism software version 9.0.0 (Boston, Massachusetts USA). This fitting process allowed for the accurate determination of IC_50_ values, providing crucial insights into the compound’s efficacy in modulating the targeted interactions or dimerization events.

### Safety

There are no hazards or risks associated with the reported work.

## ASSOCIATED CONTENT

### Supporting Information

The following files are available free of charge.

Supplementary Tables 1-7

Supplementary Figures 1-10

## AUTHOR INFORMATION

### Author Contributions

The manuscript was written through contributions of all authors. All authors have given approval to the final version of the manuscript.

## ACKNOWLEDGMENT

The authors wish to thank Zsuzsanna Izsvák and Zoltán Ivics for kindly providing components of the SB100X transposon system and Sigrid Schnögl for carefully reading the manuscript.

## ABBREVIATIONS

aa: amino acids
ATXN1: ataxin-1
cc: coiled-coil
CT: C-terminal
CNN: convolutional neural network
FL: full-length
IIBs: intranuclear inclusion bodies
MD: molecular dynamics
NT: N-terminal
Rg: radius of gyration
RMSD: root mean square deviation
RMSF: root mean square fluctuation
polyQ: polyglutamine
SCA1: Spinocerebellar ataxia type 1

## TOC graphic

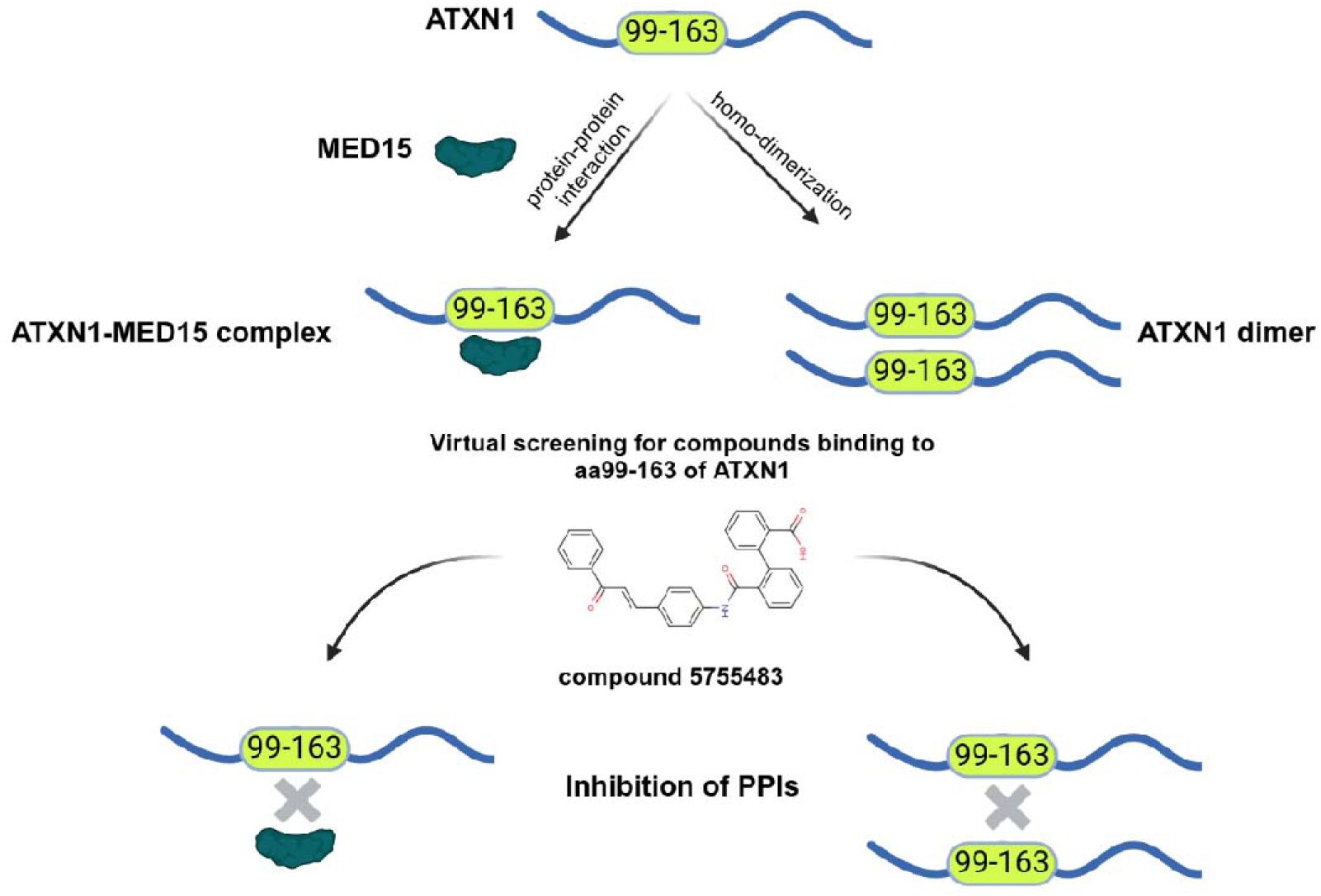

