## Supplementary Figures and Tables for "Chemical targeting of the ATXN1 aa99-163 interaction site suppresses polyQ-expanded protein dimerization"

[#] equal contribution

### Table of contents

**Supporting Tables** .......... 3
**Supporting Figures** .......... 14

### Supporting Tables

**Supplementary Table 1.** Predicted interaction sites between ATXN1 and MED15 proteins along with the simulations in which these sites were identified.

| Interaction site | Simulation |
|---|---|
| ATXN1 99-163 | $ATXN1^{NT}$/MED15 (Model 1 & 2) |
| ATXN1 198-248 | $ATXN1^{NT}$/MED15 (Model 1) |
| ATXN1 328-417 | $ATXN1^{CT}$/MED15 (Model 1) |
| ATXN1 493-540 | ATXN1(Q30)/MED15 (Model 2) |
| ATXN1 647-691 | ATXN1(Q30)/MED15 (Model 1) |
| MED15 101-294 | $ATXN1^{NT}$/MED15 (Model 1) & ATXN1(Q30)/MED15 (Model 1) |
| MED15 195-294 | $ATXN1^{NT}$/MED15 (Model 1) & ATXN1(Q30)/MED15 (Model 1) |
| MED15 395-443 | ATXN1(Q30)/MED15 (Model 2) |
| MED15 548-665 | $ATXN1^{CT}$/MED15 (Model 1) |
| MED15 635-695 | ATXN1(Q30)/MED15 (Model 1) |

**Supplementary Table 2.** Summary of MD simulation for the ATXN1-MED15 PPI. The table shows the time (in nanoseconds) of the MD simulation, the root mean square deviation (RMSD) of the protein-protein complex (in Å), the radius of gyration (Rg) (in Å) and the binding free energy (in kcal/mol) calculated over the simulation period.

| Time (ns) | RMSD (Å) | Radius of Gyration (Rg) (Å) | Binding Free Energy (kcal/mol) |
|---|---|---|---|
| 0 | 1.233390413 | 20.53983345 | -10.24775633 |
| 1 | 1.167333387 | 20.63645966 | -10.40618669 |
| 2 | 1.349256029 | 20.60584344 | -10.19841672 |
| 3 | 1.170095652 | 20.61678289 | -9.760286728 |
| 4 | 1.394108264 | 20.81145451 | -9.876092542 |
| 5 | 1.32522846 | 20.62533 | -10.22742317 |
| 6 | 1.288887956 | 20.56927624 | -10.02419018 |
| 7 | 1.292328829 | 20.63225852 | -10.10120192 |
| 8 | 1.285927571 | 20.70448506 | -10.25072222 |
| 9 | 1.243923523 | 20.75320537 | -10.15251764 |
| 10 | 1.161691807 | 20.46666841 | -10.53448288 |
| 11 | 1.166870324 | 20.45511244 | -10.95682898 |
| 12 | 1.188961143 | 20.22664954 | -10.8453488 |
| 13 | 1.14023835 | 20.27762476 | -11.00099787 |
| 14 | 1.13137454 | 20.19412859 | -11.15719778 |
| 15 | 1.085961076 | 20.38834081 | -10.73417704 |
| 16 | 1.066611404 | 20.3908357 | -10.87524656 |
| 17 | 1.110124127 | 20.38352761 | -11.02000647 |
| 18 | 1.144135442 | 20.26116532 | -10.97272047 |
| 19 | 1.160860057 | 20.52709683 | -10.60847443 |
| 20 | 1.155201131 | 20.45095285 | -10.35121857 |
| 21 | 1.256572972 | 20.65988691 | -10.6733866 |
| 22 | 1.215732922 | 20.67769528 | -10.30404328 |
| 23 | 1.2835257 | 20.6773254 | -10.39258958 |
| 24 | 1.323576293 | 20.6588815 | -9.912523616 |
| 25 | 1.230009713 | 20.74204734 | -9.905043432 |
| 26 | 1.255493396 | 20.65248874 | -10.11242282 |
| 27 | 1.444115443 | 20.83178103 | -10.08798086 |
| 28 | 1.308845568 | 20.55913411 | -9.745234221 |
| 29 | 1.246280049 | 20.62931106 | -10.37279783 |
| 30 | 1.236829172 | 20.34424823 | -10.13531703 |
| 31 | 1.085426215 | 20.49489969 | -10.56388336 |
| 32 | 1.062388493 | 20.3556512 | -11.17491644 |
| 33 | 1.08948792 | 20.17308745 | -11.21111501 |
| 34 | 1.121224795 | 20.41116159 | -11.38842832 |
| 35 | 1.047757025 | 20.2793613 | -10.89354861 |
| 36 | 1.126126869 | 20.34365811 | -10.84542025 |

| | | | |
|---|---|---|---|
| 37 | 1.179740495 | 20.35384193 | -10.8169487 |
| 38 | 1.113317333 | 20.39719901 | -10.65332235 |
| 39 | 1.196239208 | 20.57058017 | -10.41696394 |
| 40 | 1.087578267 | 20.54891044 | -10.51817066 |
| 41 | 1.226851788 | 20.5648896 | -10.40307103 |
| 42 | 1.329376684 | 20.75037584 | -10.22719866 |
| 43 | 1.302676025 | 20.7087347 | -9.944536836 |
| 44 | 1.245276346 | 20.70942462 | -9.832297624 |
| 45 | 1.238210575 | 20.57639029 | -9.871569657 |
| 46 | 1.337320445 | 20.66043841 | -10.1095434 |
| 47 | 1.329840214 | 20.76271061 | -10.31451467 |
| 48 | 1.272635771 | 20.74621218 | -10.4112137 |
| 49 | 1.189725484 | 20.57299591 | -10.4376036 |
| 50 | 1.257422176 | 20.60299783 | -10.517525 |
| 51 | 1.152711133 | 20.39739186 | -10.91269835 |
| 52 | 1.143744319 | 20.36069661 | -10.68390103 |
| 53 | 1.114129559 | 20.26836731 | -11.12946262 |
| 54 | 1.162605426 | 20.29878895 | -11.01820792 |
| 55 | 1.108440314 | 20.29729279 | -10.74278809 |
| 56 | 1.086740188 | 20.2900414 | -10.80645183 |
| 57 | 1.265122183 | 20.33748974 | -10.85313126 |
| 58 | 1.198582234 | 20.5687777 | -10.62941442 |
| 59 | 1.126820106 | 20.29096278 | -10.2816209 |
| 60 | 1.162059696 | 20.43393943 | -10.25565869 |
| 61 | 1.314554235 | 20.54068851 | -10.35980487 |
| 62 | 1.272492578 | 20.63822986 | -10.13085468 |
| 63 | 1.173515418 | 20.79874609 | -10.05905475 |
| 64 | 1.340104679 | 20.7330102 | -10.29353595 |
| 65 | 1.306290487 | 20.56100723 | -9.663966183 |
| 66 | 1.367331785 | 20.60783416 | -10.19277046 |
| 67 | 1.282563227 | 20.65634159 | -9.733883476 |
| 68 | 1.244534426 | 20.6355926 | -10.20489378 |
| 69 | 1.257024968 | 20.51128054 | -10.40452427 |
| 70 | 1.134037008 | 20.57024391 | -10.59154464 |
| 71 | 1.231818063 | 20.45572659 | -10.69574302 |
| 72 | 1.147359581 | 20.65673538 | -10.87961025 |
| 73 | 1.122417677 | 20.49243393 | -11.11207483 |
| 74 | 1.164872861 | 20.23529516 | -10.92192414 |
| 75 | 1.081756435 | 20.25914327 | -10.80926475 |
| 76 | 1.114411456 | 20.26874691 | -10.38287692 |
| 77 | 1.140600422 | 20.35075308 | -10.99664775 |
| 78 | 1.129657754 | 20.42522411 | -10.44101827 |
| 79 | 1.219329076 | 20.48803316 | -10.46985934 |

| | | | |
|---|---|---|---|
| 80 | 1.036228704 | 20.44654869 | -10.64899419 |
| 81 | 1.291711058 | 20.61038072 | -9.937401528 |
| 82 | 1.263220909 | 20.63198576 | -10.08218102 |
| 83 | 1.298333824 | 20.57573131 | -10.15360183 |
| 84 | 1.366074629 | 20.49811598 | -10.0626218 |
| 85 | 1.208266544 | 20.50517436 | -10.01281039 |
| 86 | 1.266115536 | 20.74489559 | -10.47932158 |
| 87 | 1.255879782 | 20.70344547 | -10.23938602 |
| 88 | 1.2708134 | 20.53134962 | -10.28361045 |
| 89 | 1.149864478 | 20.55776191 | -10.66644876 |
| 90 | 1.234720942 | 20.58681897 | -10.23557236 |
| 91 | 1.203583458 | 20.40986473 | -11.14980578 |
| 92 | 1.026966693 | 20.3581714 | -10.96583487 |
| 93 | 1.180180648 | 20.25006208 | -10.94919881 |
| 94 | 1.101495868 | 20.34322453 | -10.82430066 |
| 95 | 1.08574593 | 20.42357565 | -10.61826189 |
| 96 | 1.214489229 | 20.31650363 | -11.15921909 |
| 97 | 1.093182606 | 20.28198517 | -10.88471586 |
| 98 | 1.197612107 | 20.40301551 | -10.68797643 |
| 99 | 1.095876053 | 20.49306131 | -10.31052242 |
| 100 | 1.224798891 | 20.62646648 | -10.49975704 |
| **Median** | 1.198582234 | 20.53134962 | -10.4376036 |

**Supplementary Table 3.** Aggregation-prone residues and their corresponding values. The table presents ATXN1 residues (aa90-170) along with their average aggregation-propensity values per amino acid (a4v) and the HSA (Hot-Spot Area) for each aa residue which define the aggregation profile (AP) of the polypeptide based on its complete sequence. Aminoacids were ranked according to their HSA value.

| aa | a4v | HSA |
|---|---|---|
| H125 | 0.368 | 2.151 |
| T126 | 0.214 | 2.151 |
| F127 | 0.19 | 2.151 |
| Q128 | 0.257 | 2.151 |
| F129 | 0.02 | 2.151 |
| I130 | 0.156 | 2.151 |
| G131 | 0.223 | 2.151 |
| S132 | 0.189 | 2.151 |
| S133 | 0.015 | 2.151 |
| Q134 | 0.232 | 2.151 |
| Y135 | 0.069 | 2.151 |
| G137 | 0.085 | 2.079 |
| T138 | 0.277 | 2.079 |
| Y139 | 0.274 | 2.079 |
| A140 | 0.359 | 2.079 |
| S141 | 0.142 | 2.079 |
| F142 | 0.294 | 2.079 |
| I143 | 0.508 | 2.079 |
| V98 | 0.153 | 1.415 |
| A99 | 0.18 | 1.415 |
| T100 | 0.289 | 1.415 |
| T101 | 0.421 | 1.415 |
| L102 | 0.273 | 1.415 |
| S145 | 0.356 | 1.241 |
| Q146 | 0.345 | 1.241 |
| L147 | 0.369 | 1.241 |
| I148 | 0.091 | 1.241 |
| V155 | 0.045 | 0.67 |
| T156 | 0.049 | 0.67 |
| S157 | 0.06 | 0.67 |
| A158 | 0.06 | 0.67 |
| V159 | 0.13 | 0.67 |
| A160 | 0.157 | 0.67 |
| S161 | -0.002 | 0.67 |
| A162 | -0.002 | 0.67 |
| A163 | -0.006 | 0.67 |

| | | |
|---|---|---|
| V118 | 0.21 | 0.594 |
| Q119 | 0.194 | 0.594 |
| Y120 | 0.13 | 0.594 |
| T151 | 0.082 | 0.349 |
| A152 | 0.168 | 0.349 |
| N153 | 0.039 | 0.349 |
| H122 | 0.157 | 0.272 |
| L123 | 0.075 | 0.272 |
| A104 | 0.114 | 0.168 |
| A105 | 0.005 | 0.168 |
| Y106 | -0.011 | 0.168 |
| V115 | -0.01 | 0.13 |
| S116 | 0.099 | 0.13 |
| R94 | -0.004 | 0.095 |
| S95 | 0.012 | 0.095 |
| V96 | 0.027 | 0.095 |
| S91 | -0.204 | 0 |
| A92 | -0.067 | 0 |
| P93 | -0.028 | 0 |
| P97 | 0.18 | 0 |
| P103 | 0.289 | 0 |
| A107 | -0.045 | 0 |
| T108 | -0.185 | 0 |
| P109 | -0.185 | 0 |
| Q110 | -0.037 | 0 |
| P111 | -0.06 | 0 |
| G112 | -0.196 | 0 |
| T113 | -0.048 | 0 |
| P114 | -0.145 | 0 |
| P117 | 0.036 | 0 |
| A121 | -0.029 | 0 |
| P124 | 0.09 | 0 |
| S136 | -0.123 | 0 |
| P144 | 0.492 | 0 |
| P149 | -0.105 | 0 |
| P150 | 0.07 | 0 |
| P154 | 0.018 | 0 |
| G164 | -0.03 | 0 |
| A165 | -0.286 | 0 |
| T166 | -0.396 | 0 |
| T167 | -0.396 | 0 |
| P168 | -0.504 | 0 |
| S169 | -0.376 | 0 |

| | | |
|---|---|---|
| Q170 | -0.455 | 0 |

**Supplementary Table 4.** Prediction and evaluation of binding pockets in the ATXN1$^{NT}$ protein identified using the PockDrug online server. The table indicate volume hull, hydrophobic kyte, polar and aromatic residues proportion, frequency of Otyr atoms, number of pocket residues, druggability probability score and standard deviation.

| Pocket | Volume Hull | Hydrophobic kyte | Polar residues proportion | Aromatic residues proportion | Otyr atom | Number of pocket residues | Druggability Probability | STDEV |
|---|---|---|---|---|---|---|---|---|
| **1** | 1193.85 | -0.05 | 0.4 | 0.13 | 0.02 | 15 | 0.81 | 0.04 |
| **2** | 796.71 | -0.34 | 0.56 | 0.25 | 0 | 16 | 0.74 | 0.1 |
| **3** | 3436.6 | -0.43 | 0.49 | 0.14 | 0.01 | 35 | 0.71 | 0.02 |

**Supplementary Table 5.** Binding score of the selected and AI-predicted compounds (n=24) with high affinity for ATXN1 aa99-163.

| ChemBridge code | IUPAC NAME | Score |
|---|---|---|
| 6418951 | 2-{[4-(3,4-dihydro-2(1H)-isoquinolinylmethyl)benzoyl]amino}benzoic acid | 0.979 |
| 5466832 | 3-(aminocarbonyl)-1-[2-(4-biphenylyl)-1-methyl-2-oxoethyl]pyridinium bromide | 0.974 |
| 6015799 | ethyl 4-{5-[({3-[(2-methylbenzoyl)amino]phenyl}amino)carbonyl]-2-furyl}benzoate | 0.975 |
| 5759625 | 2-[3-(1,3-benzoxazol-2-yl)phenyl]-1,3-dioxo-5-isoindolinecarboxylic acid | 0.989 |
| 5922064 | 3-[(3-{[(4-phenoxyphenyl)amino]sulfonyl}benzoyl)amino]benzoic acid | 0.968 |
| 5871031 | 4-(benzyloxy)-N-(3-[1,3]oxazolo[4,5-b]pyridin-2-ylphenyl)benzamide | 0.982 |
| 573393 | 1,3-dioxo-2-[4-(5-phenyl-1,3,4-oxadiazol-2-yl)phenyl]-5-isoindolinecarboxylic acid | 0.975 |
| 7629383 | N-3-quinolinyl-9H-fluorene-2-sulfonamide | 0.989 |
| 7779825 | (4-{[(2-benzyl-1,3-dioxo-2,3-dihydro-1H-isoindol-5-yl)carbonyl]amino}phenoxy)acetic acid | 0.975 |
| 7933413 | N-[3-(2-quinoxalinyl)phenyl]benzamide | 0.984 |
| 6433773 | 5-[(2-methylbenzoyl)amino]-N-(3-methylphenyl)-1-phenyl-1H-pyrazole-4-carboxamide | 0.976 |
| 5751467 | N-(2-methyl-5-[1,3]oxazolo[4,5-b]pyridin-2-ylphenyl)-2-naphthamide | 0.972 |
| 5160674 | 2-dibenzo[b,d]furan-3-yl-1,3-dioxo-5-isoindolinecarboxylic acid | 0.986 |
| 6238693 | 3'-[(9H-fluoren-2-ylcarbonyl)amino]-4-biphenylyl acetate | 0.99 |
| 6048589 | 4-{4-[(diphenylacetyl)amino]benzoyl}phthalic acid | 0.993 |
| 6052953 | 2-({[4-(2-naphthyloxy)phenyl]amino}carbonyl)benzoic acid | 0.984 |
| 7969487 | N-{3-[(1,3-dioxo-2,3-dihydro-1H-isoindol-5-yl)oxy]phenyl}-2-phenylbutanamide | 0.923 |
| 6846586 | 4-tert-butyl-N-[3-(5-phenyl-1,3,4-oxadiazol-2-yl)phenyl]benzamide | 0.985 |
| 5561416 | 1-(4-biphenylyl)-2-{2-imino-3-[2-(4-morpholinyl)ethyl]-2,3-dihydro-1H-benzimidazol-1-yl}ethanone hydrobromide | 0.964 |
| 5606311 | N-1-naphthyl-N'-{3-[2-(2-pyridinyl)ethyl]phenyl}urea | 0.979 |
| 5319422 | 3-(benzoylamino)-1-[2-(9H-fluoren-2-yl)-2-oxoethyl]pyridinium bromide | 0.981 |
| 6032246 | 3-(benzyloxy)-N-(2-methyl-3-[1,3]oxazolo[4,5-b]pyridin-2-ylphenyl)benzamide | 0.958 |
| 5780637 | cyclohexyl (4-methyl-3-{[(1-naphthylamino)carbonyl]amino}phenyl)carbamate | 0.988 |
| 5755483 | 2'-({[4-(3-oxo-3-phenyl-1-propen-1-yl)phenyl]amino}carbonyl)-2-biphenylcarboxylic acid | 0.967 |

**Supplementary Table 6.** Key residues between ATXN1 aa99-163 and compound 5755483 identified through molecular docking and MD simulations. The table lists the interacting residues, the interaction type, the frequency (%) of the interaction during the simulation, the mean interaction distance (nm) and the binding free energy (kcal/mol).

| Residue | Interaction Type | Interaction Frequency (%) | Mean Distance (nm) | Binding free energy (kcal/mol) |
|---|---|---|---|---|
| $Thr^{113}$ | Hydrophobic | 65 | 0.2 | -2.5 |
| $Pro^{114}$ | Hydrophobic | 65 | 0.24 | -2.2 |
| $Tyr^{135}$ | Pi-Pi stacking | 67 | 0.35 | -4.1 |
| $Ser^{136}$ | Hydrogen bond | 85 | 0.3 | -5.8 |

**Supplementary Table 7.** Primer sequences used for the generation of deletion-carrying ATXN1 and MED15 Gateway entry clones.

| Deletion clone | Forward Primer (5’-3’) | Reverse Primer (5’-3’) |
|---|---|---|
| ATXN1 Δ99-163 | GGGGCCACCACTCCATC | CACGGGGACAGACCTGGG |
| ATXN1 Δ198-248 | CAGTACGTCCACATTTCCAGTTCT | CTGCTCAGCCTTGTGTCCCGG |
| ATXN1 Δ328-417 | GGCCTGCATTTAGGGAAGC | CAGGACCTCCTTGGCCTG |
| ATXN1 Δ647-691 | AACCTGAAGAACGGCTCTGTT | TTCAACGCTGACCTGGGC |
| ATXN1 Δ493-540 | CTGGTCACCCAGGCCGCC | GGGCTGTGTGCCGGGGATC |
| MED15 Δ101-294 | CACACACAGCACCACCAG | GCCAATTCCAGCGGCTCC |
| MED15 Δ635-695 | TGGATAGACCGGCAGTGGCAG | GATGCTCTGCCGCTCATCATC |
| MED15 ΔCC | CCATCACAACTCCCGCCACAGTCGC | TCCCGCAGCAGGTCCGCCAGTCAGG |
| MED15 Δ195-294 | CACACACAGCACCACCAG | ACTCTGCTGAGCCTGGAACT |
| MED15 Δ395-443 | CCCTCACCGCAGCCCTCC | CGAGGACAGCATGGGGAGG |
| MED15 Δ548-665 | CTGATCTGCAAGCTGGATGACA | CTTTTGCAAGGTCTTCAGGGG |
| ATXN1 ΔAXH | CTTACCCTCAAGAACCTGAAGAACG | CACAGGCAGGTGGATCTGGGCCTGCA |

### Supporting Figures

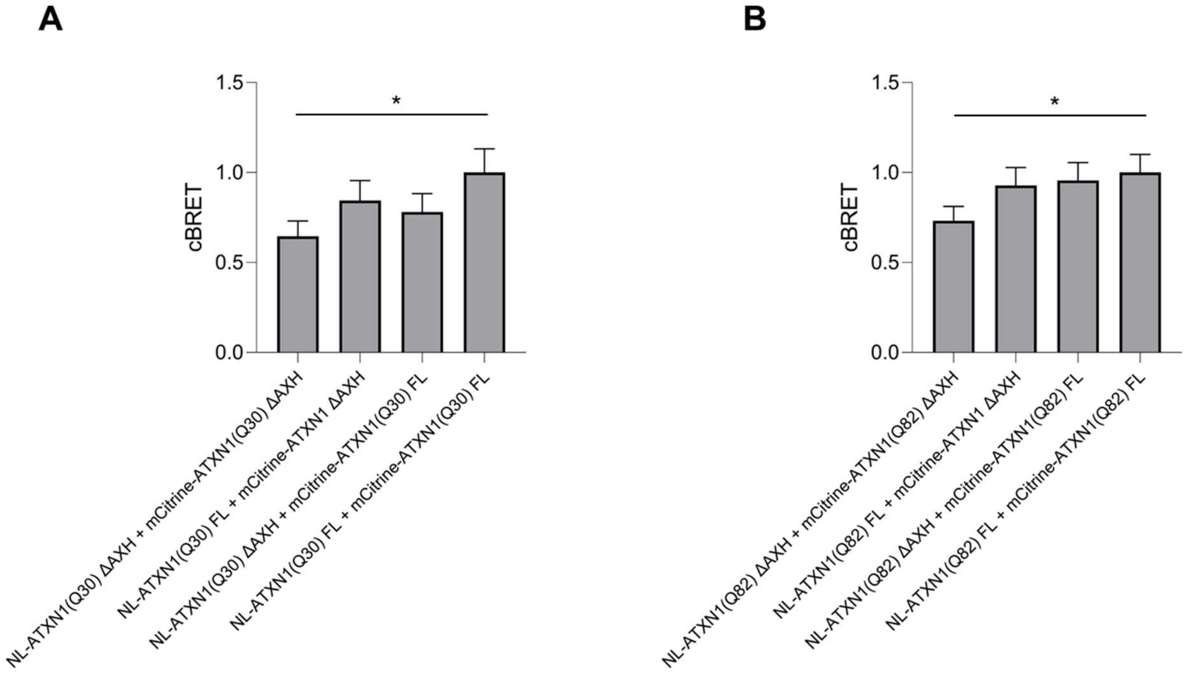


**Supplementary Figure 1.** Quantification of **(A)** wild-type (Q30) and **(B)** polyQ-expanded (Q82) ATXN1 homo-dimerization harboring an AXH domain deletion. Deletion of the AXH domain (ΔAXH) moderately reduces ATXN1(Q30) or (Q82) dimerization compared to the respective full-length (FL) proteins. Data are presented as mean ± standard deviation (SD).

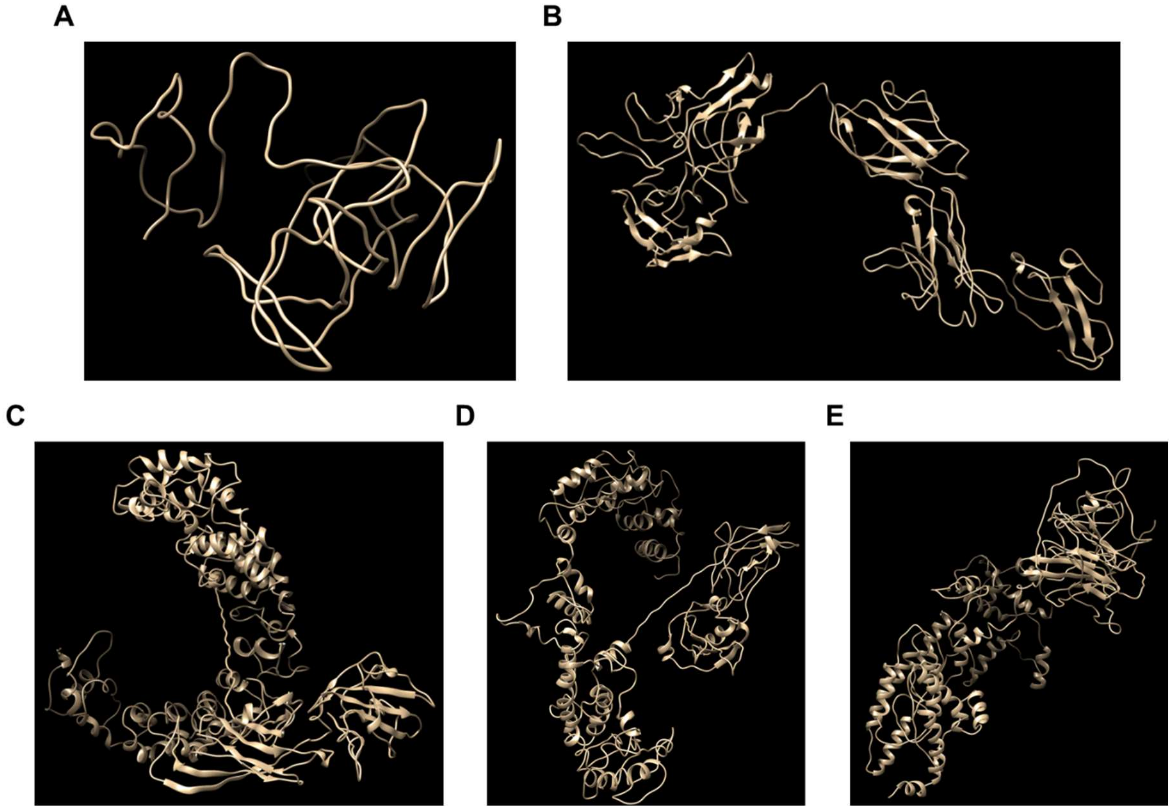


**Supplementary Figure 2**. Predicted structures of ATXN1 or MED15 proteins using the I-TASSER software. **(A)** ATXN1$^{NT}$, **(B)** ATXN1$^{CT}$, **(C)** ATXN1(Q30), **(D)** MED15 (model 1) and **(E)** MED15 (model 2)

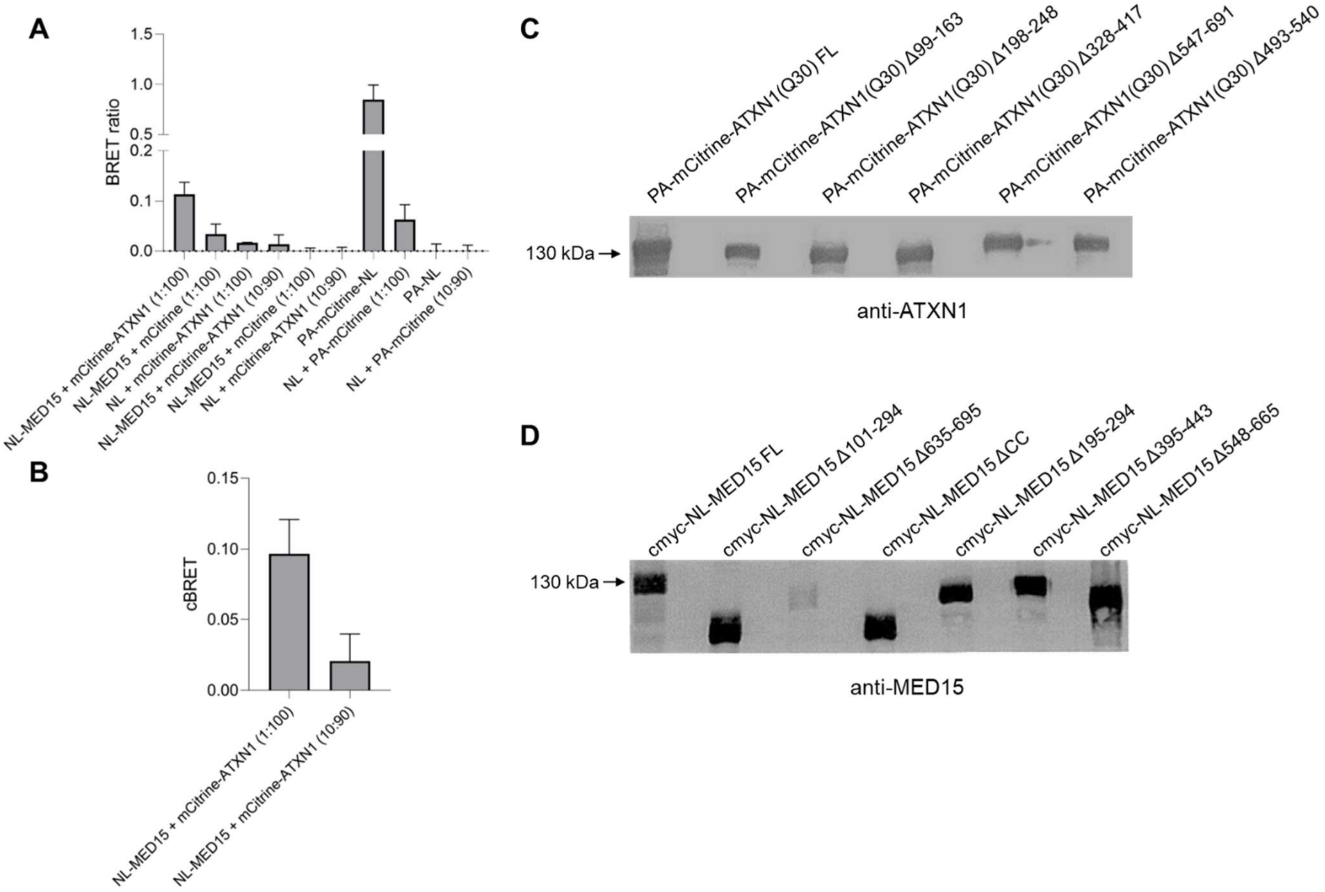


**Supplementary Figure 3.** Quantification of ATXN1-MED15 PPI using LuTHy assay. **(A)** BRET ratios for ATXN1-MED15 PPI. The proteins PA-mCitrine-ATXN1 and NL-MED15 (at a 1:100 ratio), along with the positive control PA-mCitrine-NL, show significantly elevated BRET ratios, indicating a strong interaction. **(B)** cBRET ratios confirming the efficient quantification of ATXN1-MED15 PPI at a 1:100 ratio of PA-mCitrine-ATXN1/NL-MED15. **(C)** Immunoblot for ATXN1 in extracts from transiently transfected HEK293T producing full-length or deletion-carrying ATXN1 protein. **(D)** Immunoblot for MED15 in extracts from transiently transfected HEK293T producing full-length or deletion-carrying MED protein.

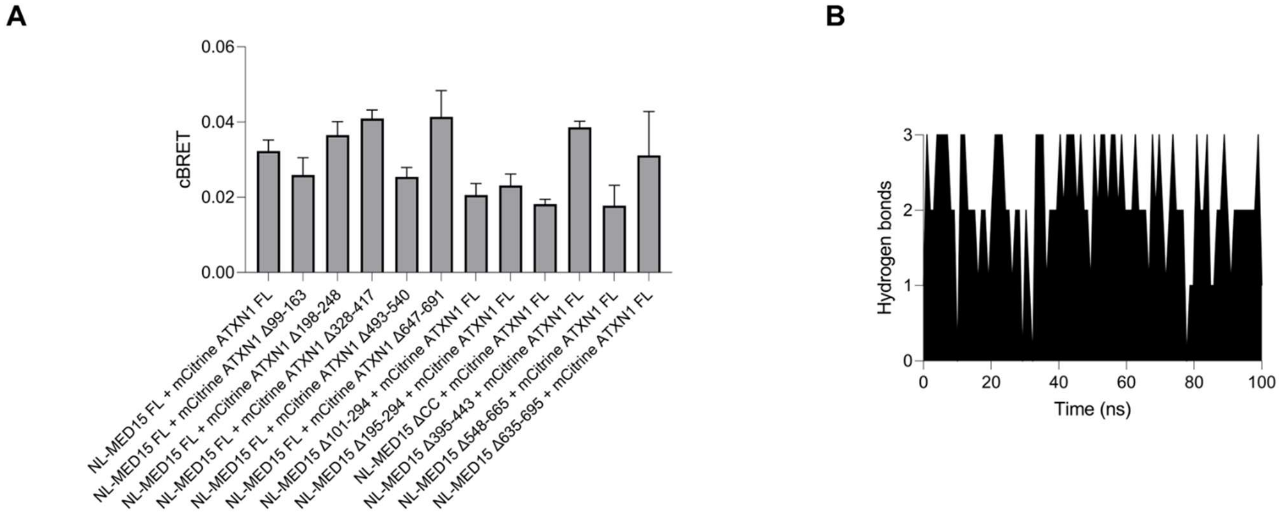


**Supplementary Figure 4. (A)** Quantification (cBRET ratio) of the interaction between full-length and deletion-carrying ATXN1/MED15 proteins. Results were compared to the interaction of full-length ATXN1 and MED15. **(B)** The number of hydrogen bonds formed between domains 99-163 of ATXN1 and 548-665 of MED15 over a 100 ns simulation.

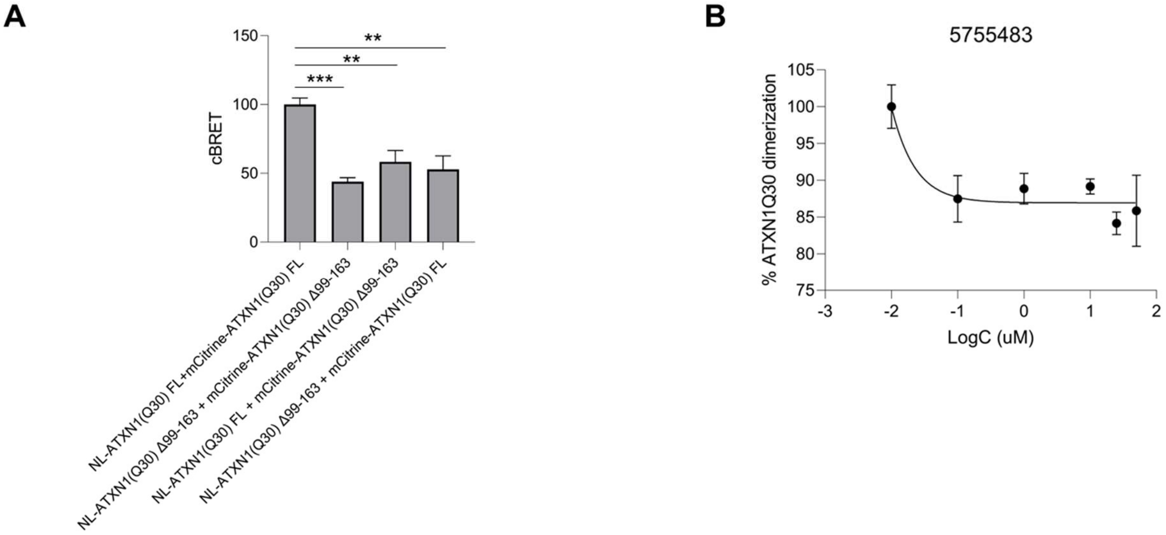


**Supplementary Figure 5.** The aa99-163 domain mediates the homo-dimerization of wild-type ATXN1 in cell models. **(A)** Quantification of the homo-dimerization of full-length or Δ99-163 ATXN1(Q30) using the LuTHy assay. The bar graph shows various combinations of full-length or deletion-carrying ATXN1 proteins. Data in bar graphs are shown as mean ± SD. ** p-value <0.01, *** p-value < 0.001). **(B)** Dose-dependent effect of compound 5755483 in the homo-dimerization of ATXN1(Q30).

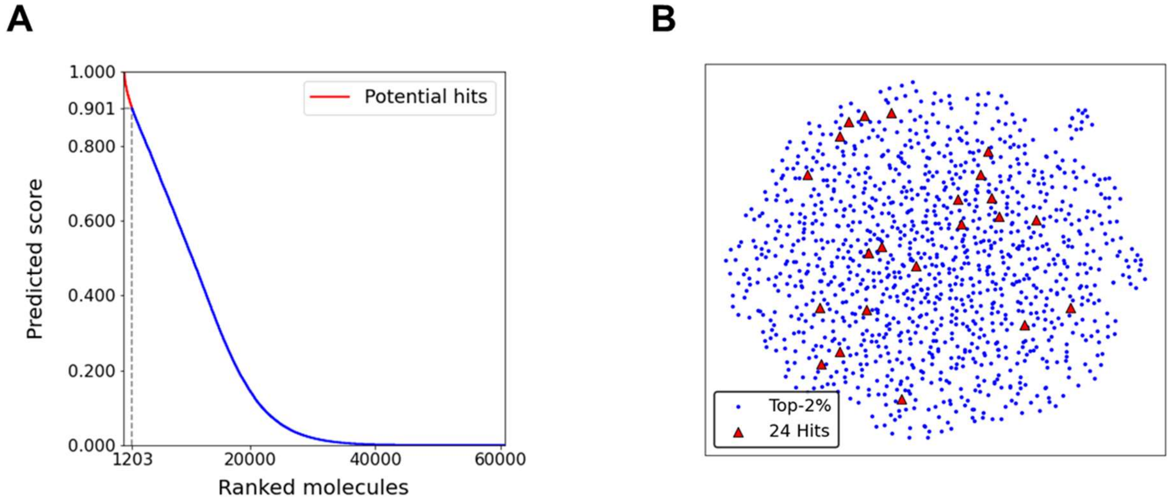


**Supplementary Figure 6. (A)** Molecule ranking scores after applying the virtual screening pipeline to the initially filtered library of approximately 60,000 molecules, followed by the selection of the top 2% (1,203 molecules). **(B)** UMAP visualization of the final selected hits (n=24) compared to the top 2% of screened molecules, projected onto a reduced feature space defined by shape descriptors. This visualization highlights the distribution and clustering of the final hits relative to the broader set of top-ranked molecules.

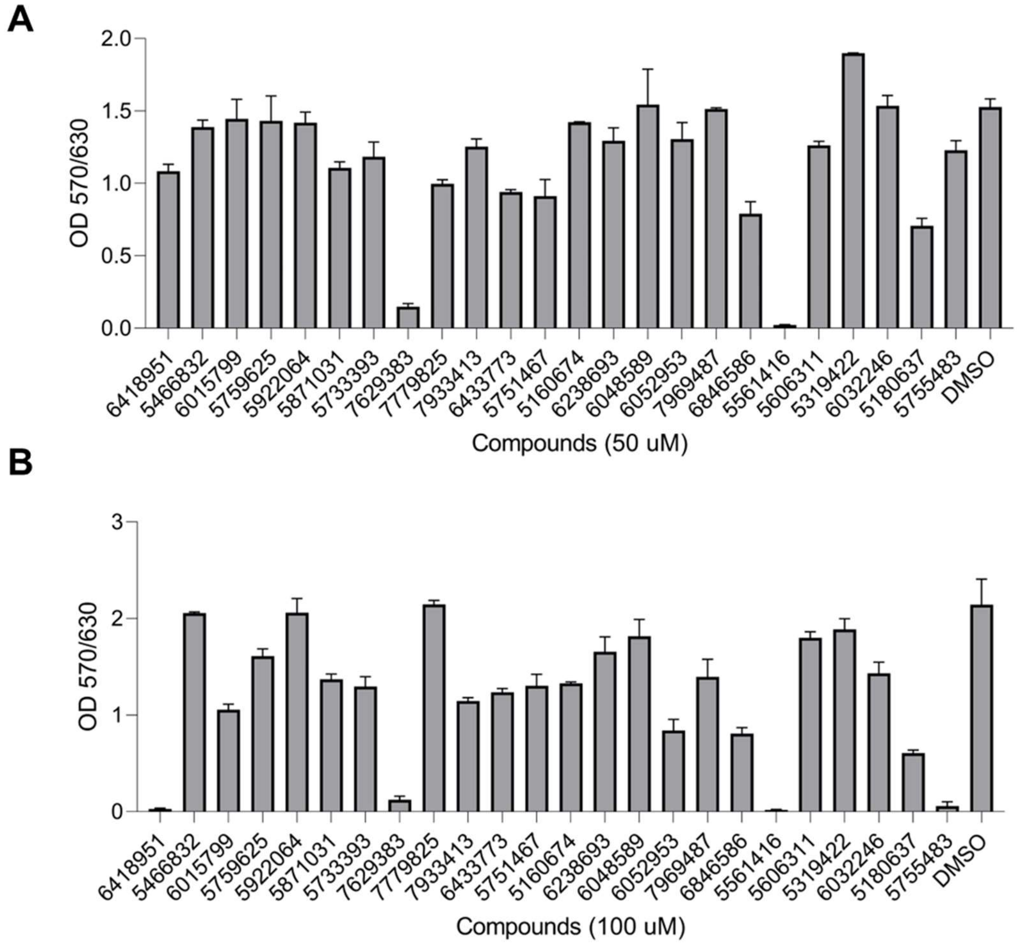


**Supplementary Figure 7.** Viability of HEK293T cells following a 48-hour treatment with compounds at concentrations of **(A)** 100 μM and **(B)** 50 μM, compared to control treatment with DMSO.

6032246

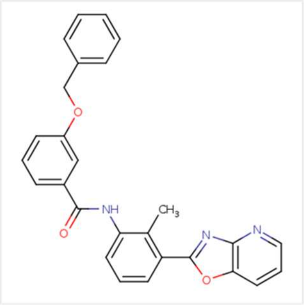

6846586

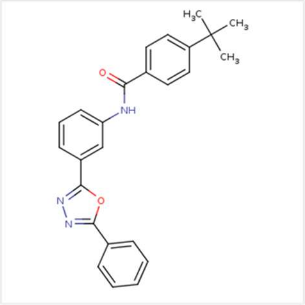

5751467

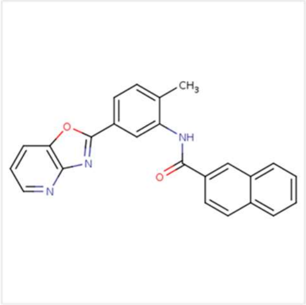

5319422

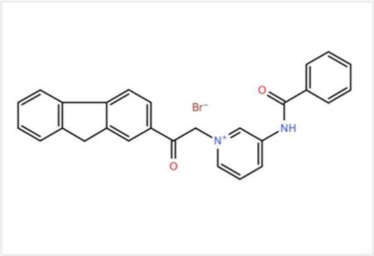

7969487

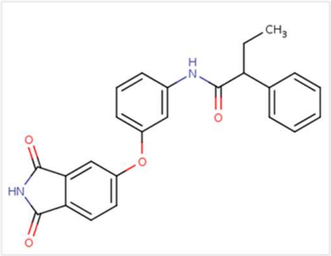

**Supplementary Figure 8.** Chemical structures of five compounds (Chembridge ID: 6032246, 6846586, 5751467, 5319422, 7969487) reducing ATXN1-MED15 PPI.

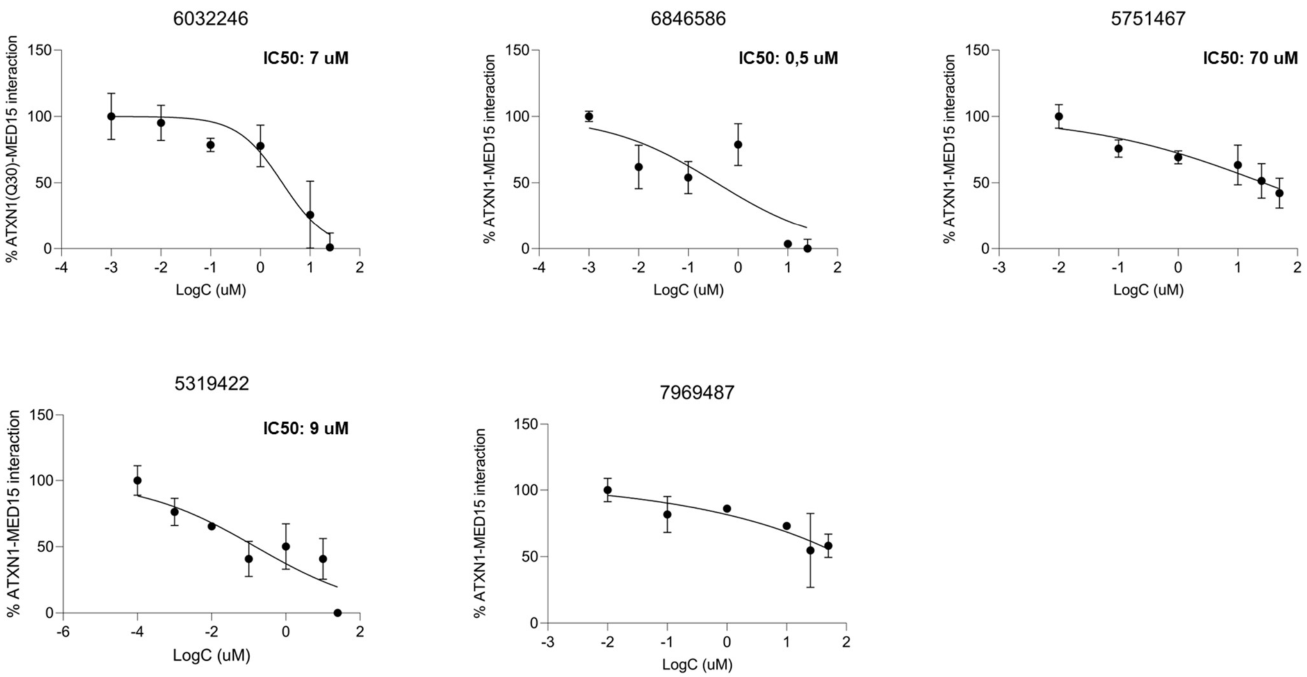


**Supplementary Figure 9.** Concentration-response curves for the inhibitory effect of selected compounds on ATXN1-MED15 PPI.

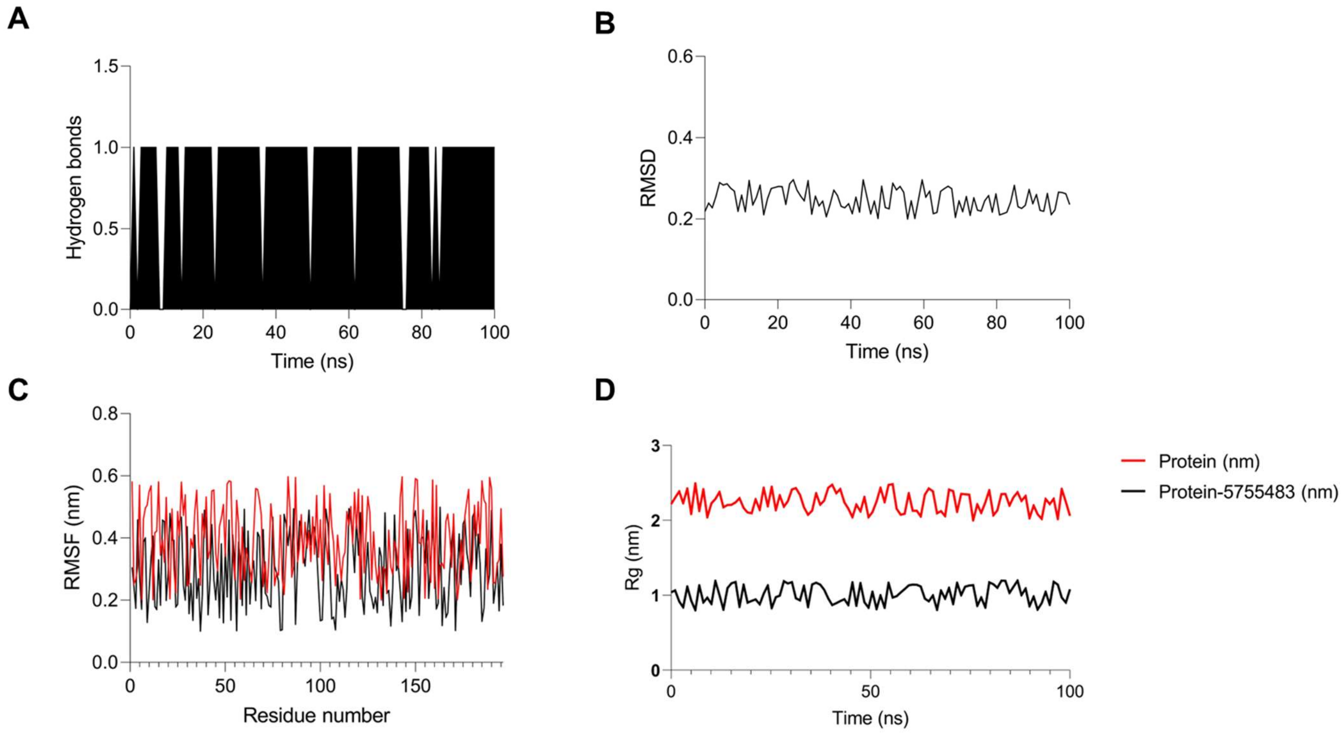


**Supplementary Figure 10.** MD simulation for the interaction between ATXN1 aa99-163 and compound 5755483. **(A)** The number of hydrogen bonds formed between ATXN1 and 5755483 over 100 ns of simulation. **(B)** RMSD of the ATXN1-5755483 complex over the simulation time. **(C)** RMSF and **(D)** Radius of gyration (Rg) of the ATXN1 protein in the presence (black) or absence (red) of compound 5755483. The lower Rg values of the bound state indicate that compound 5755483 promotes a more stable protein conformation.
